# Covalent Modulators of Immune Regulatory Transcription Factors IRF8 and IRF5

**DOI:** 10.1101/2025.08.03.668300

**Authors:** Thang Cong Do, Henry Chan, Justin Greene, Sam Sabaat, Connor Ludwig, Nathan S. Abell, Matthew L. Albert, Sriram Kosuri, Daniel K. Nomura

## Abstract

Transcription factors are among the most challenging targets for drug discovery due to their lack of classical binding pockets and high degree of intrinsic disorder, despite the therapeutic importance of many of these proteins. IRF5 and IRF8 are key transcriptional regulators of innate immune signaling that orchestrate pro-inflammatory gene expression programs in response to stimuli such as toll-like receptor activation, making them central players in autoimmune and inflammatory diseases. Despite their therapeutic interest, direct targeting of IRF5 and IRF8 has remained challenging. Through a screen originally intended to identify cysteine-reactive electrophilic ligands that could directly and covalently engage and degrade IRF5, we identified an acrylamide hit, EN1033, that not only targeted and degraded IRF5 but also degraded a related inflammatory transcription factor, IRF8, more robustly and rapidly. EN1033 destabilized and degraded IRF5 and IRF8 by covalently targeting C28 and C223, respectively, as evidenced by the attenuation of their degradation through mutagenesis of these cysteines. We also found that IRF8 loss led to downregulation and inhibition of IRF5 activity, suggesting crosstalk between these two transcription factors, both of which are targeted by EN1033. Upon exploring structure-activity relationships, we identified an optimized compound, TH-B10, that more potently and selectively targeted and degraded IRF8 and, secondarily, downregulated IRF5 by targeting C223 on IRF8. Overall, we identify an early-stage pathfinder molecule, TH-B10, that directly covalently targets and degrades IRF8 and secondarily modulates IRF5 to shut down pro-inflammatory transcriptional activity.

## Introduction

Among undruggable proteins, transcription factors remain some of the most challenging targets for ligand discovery due to their lack of classical binding pockets, shallow protein-protein interaction interfaces, and high degree of intrinsic disorder ^1,2^. In immunology, the interferon regulatory factor (IRF) class of transcription factors plays central roles in coordinating innate and adaptive immune responses, particularly through the regulation of interferon signaling and inflammatory gene expression ^3,4^. Consisting of nine mammalian family members (IRF1-IRF9), these factors are activated downstream of pattern recognition receptors, such as toll-like and RIG-1-like receptors, and drive content-dependent transcription programs that control antiviral defense, cytokine production, antigen presentation, and immune cell differentiation. Dysregulation of IRF activity is implicated in a broad spectrum of human diseases, including autoimmune disorders, chronic inflammation, cancer, and infectious diseases, positioning IRFs as critical nodes in immune homeostasis and attractive targets for therapeutic intervention in inflammation and cancer ^3–7^.

Among them, IRF5 and IRF8 are particularly critical. IRF5 is a potent driver of pro-inflammatory macrophage activation and cytokine production, while IRF8 governs the development and function of monocytes, dendritic cells, and other myeloid lineages ^3,5–7^. Their pivotal roles in controlling immune cell fate and inflammatory signaling make IRF5 and IRF8 compelling therapeutic targets in immune-mediated diseases. Chronic IRF5 and IRF8 activity may drive inflammation-associated cancers, such as colon cancers, and IRF8 may drive the malignancy of select myeloid malignancies ^8–10^. While there are now reports of an IRF5 degrader from Kymera and an allosteric inhibitor of IRF5 from Hotspot Therapeutics ^11^, their structures have not yet been disclosed. To our knowledge, there are currently no small-molecule inhibitors or degraders reported for IRF8. Identifying potential approaches for ligandability and ligand discovery against these targets could reveal alternative strategies for therapeutically targeting these transcription factors.

Activity-based protein profiling (ABPP) and covalent chemoproteomic approaches have arisen as powerful approaches for targeting cryptic, allosteric, shallow, or structurally disordered sites within classically intractable disease targets, including transcription factors ^12–25^. Our lab has reported on direct-acting covalent destabilizing degraders that target intrinsically disordered cysteines within oncogenic transcription factors, including MYC, CTNNB1, and AR-V7 ^21–23^. The Cravatt lab has recently reported on a covalent ligand that stereoselectively engages a cysteine on the pioneering transcription factor FOXA1 to rewire its transcriptional activity ^16^.

In this study, we sought to identify a direct-acting, covalent degrader against the inflammatory transcription factor IRF5. During characterization, we opportunistically identified an early-stage pathfinder dual IRF5 and IRF8 covalent destabilizing degrader that preferentially acts through the covalent targeting of IRF8.

## Results

### Screening for a Covalent IRF5 Degrader

To identify covalent IRF5 degraders, we screened our cysteine-reactive covalent ligand library in THP1 macrophage leukemia cancer cells overexpressing HiBiT-tagged IRF5. While we did not find compounds that showed a loss of greater than 50% of Hibit-IRF5, possibly due to the overexpression of Hibit-IRF5, we identified acrylamide EN1033 as the top hit **(Figure 1a-1b; Table S1)**. We next demonstrated that the loss of HiBiT-IRF5 was dependent on the ubiquitin-proteasome pathway, but not the autophagy or lysosomal pathway, since we could observe complete rescue of HiBiT-IRF5 loss with both an E1 ubiquitin activating enzyme inhibitor, TAK243, and a proteasome inhibitor, bortezomib (BTZ), but not with a lysosomal v-ATPase inhibitor, bafilomycin **(Figure 1c)**. This reduction in HiBiT-IRF5 is dose-dependent, with maximal degradation of 68% at the highest concentration tested **(Figure 1d)**. We further confirmed that endogenous IRF5 levels were also reduced in unmodified THP1 cells **(Figure 1e)**. Given the relatively high concentration required for IRF5 degradation, we confirmed that EN1033 does not exhibit cytotoxicity in THP1 cells at concentrations up to 100 μM, the highest used in this study **(Figure S1a)**. While previous studies have also shown that electrophilic molecules can cause stress granule formation at high concentrations^26^, EN1033 does not induce stress granules, as observed in cells treated with the positive control, sodium arsenite **(Figure S1b)**.

**Figure 1.**
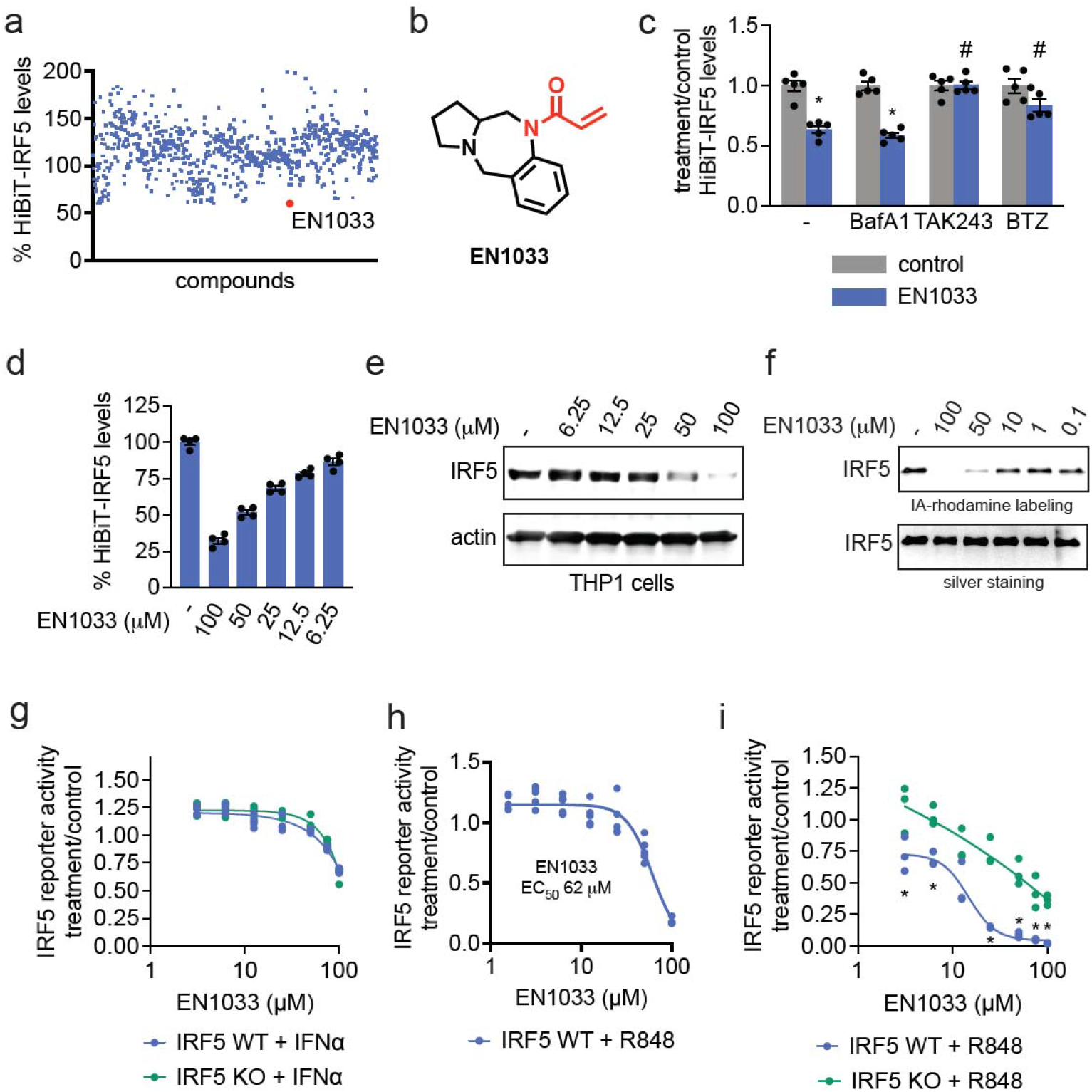
Screening for a Covalent IRF5 Degrader. **(a)** Covalent ligand screen in THP1 cells overexpressing a HiBiT-tagged IRF5. HiBiT-IRF5-expressing THP1 cells were treated with DMSO vehicle or covalent ligand (50 μM) for 24 h, after which HiBiT-IRF5 was detected by luminescence through detection with LgBiT. Individual compounds, structures, and data points are shown in **Table S1. (b)** Structure of top hit EN1033. Cysteine-reactive acrylamide warhead highlighted in red. **(c)** Ubiquitin and proteasome-dependence of HiBiT-IRF5 loss. HiBiT-IRF5-expressing THP1 cells were pre-treated with DMSO vehicle or lysosomal v-ATPase inhibitor Bafilomycin A (BafA), E1 ubiquitin conjugating enzyme inhibitor TAK243, and proteasome inhibitor BTZ (100 nM) for 1 h prior to treatment with DMSO vehicle or EN1033 (50 μM) for 24 h, after which HiBiT-IRF5 was detected by luminescence through detection with LgBiT. **(d)** Dose-response of HiBiT-IRF5 loss. HiBiT-IRF5-expressing THP1 cells were treated with DMSO vehicle or EN1033 for 24 h, after which HiBiT-IRF5 was detected by luminescence through detection with LgBiT. **(e)** Dose-response of IRF5 loss. THP1 cells were treated with DMSO vehicle or EN1033 for 24 h after which IRF5 and loading control actin levels were assessed by SDS/PAGE and Western blotting. **(f)** Gel-based ABPP of EN1033 against pure IRF5 protein. IRF5 protein was pre-incubated with DMSO vehicle or EN1033 for 1 h prior to treatment with IA-rhodamine (100 nM) for 30 min. IA-rhodamine labeling was assessed by SDS/PAGE and in-gel fluorescence and loading was assessed by silver staining. **(g, h,i)** IRF5 luciferase reporter activity. IRF5 wildtype (WT) or knockout (KO) THP1 cells expressing an IRF5 luciferase reporter were stimulated with IFNα (1000 U/mL) **(g)** or R848 (10 µM) **(h,i)** and treated with DMSO vehicle or EN1033 for 12 h **(h)** 24 h **(g, i)**, after which luciferase activity was assessed. Data in **(c,d,g,h,i)** are from n=3-4 biologically independent replicates per group. Graphs in **(c,d)** show individual replicate values and average ± sem. Blots and gels in **(e,f)** are representative of n=3 biologically independent replicates per group. Significance expressed as *p<0.05 compared to vehicle-treated controls in and compared to EN1033 treatment in THP1 KO cells in **(h)** and #p<0.05 compared to cells treated with EN1033 alone.

The electrophilic handle was necessary for this IRF5 loss, as the non-reactive analog TH3-116 did not show loss of IRF5 in THP1 cells **(Figure S2a-S2b)**. We also demonstrated that EN1033 directly binds to IRF5 *in vitro* using purified human IRF5 protein, showing that EN1033 could outcompete cysteine-reactive probe labeling in a dose-responsive manner using gel-based ABPP approaches **(Figure 1f)** ^24,25^. Through mass spectrometry (MS/MS) analysis, we also found that EN1033 covalently labels two cysteines on IRF5—C28 and C121 **(Figure S3)**. These data indicated that EN1033 directly and covalently binds to IRF5.

We next assessed the effect of EN1033 on IRF5 transcriptional activation in THP1 cells. EN1033 modestly inhibits I-SRE reporter activity in THP1 cells, whether unstimulated or stimulated with a control non-IRF5 dependent agonist IFNα. These inhibitory effects were not IRF5-dependent, showing similar activity profiles in IRF5 WT or knockout (KO) backgrounds (**Figure 1g; Figure S4)**. The unreactive control TH3-116 also did not exhibit inhibitory activity, either in the absence of an agonist or with an IRF5 pathway TLR7/8 agonist, R848 **(Figure S4)**. In contrast, we observed dose-responsive inhibition of IRF5-dependent R848-stimulated IRF5 transcriptional reporter activity, which was significantly attenuated in the THP1 IRF5 KO background, with a 50% effective concentration (EC_50_) of 62 μM at 12 h **(Figure 1h)** and 15 µM at 24 h **(Figure 1i)**. These data collectively demonstrated on-pathway inhibition of IRF5 transcriptional activity in response to EN1033 treatment. Given the high degree of IRF5 inhibition relative to the loss of protein abundance, it is possible that EN1033 binding to IRF5 both destabilizes and directly inhibits the function of IRF5.

### Validation of EN1033 as an IRF5 Degrader and Discovery of Effects on IRF8

To assess the proteome-wide selectivity of IRF5 degradation, we performed quantitative proteomic profiling on THP1 cells treated with EN1033 **(Figure 2a-2b; Table S2)**. Surprisingly, in the initial 15 h EN1033 treatment condition, among the >3,600 proteins quantified, we observed the selective loss of related IRF family member IRF8 and three other proteins, with no significant change to IRF5. With 24 h EN1033 treatment, we observed less selective, but now significant, loss of IRF5 and IRF8, along with 145 other proteins showing significant changes in levels among the >2,600 proteins quantified. These results suggested that EN1033 may be more acutely affecting IRF8 levels compared to IRF5. We confirmed IRF8 loss in THP1 cells following EN1033 treatment and demonstrated that this loss was proteasome-dependent, as evidenced by the attenuation of IRF8 loss with a proteasome inhibitor **(Figure 2c)**. We also demonstrate a dose-responsive loss of both IRF8 and IRF5 with EN1033 in THP1 cells after 24 hours of treatment following R848 stimulation **(Figure 2d)**. We also observed time-dependent reductions in both IRF8 and IRF5, with degradation starting at 8 hours of treatment **(Figure 2e)**. These results indicate that EN1033 degraded both IRF8 and IRF5.

**Figure 2.**
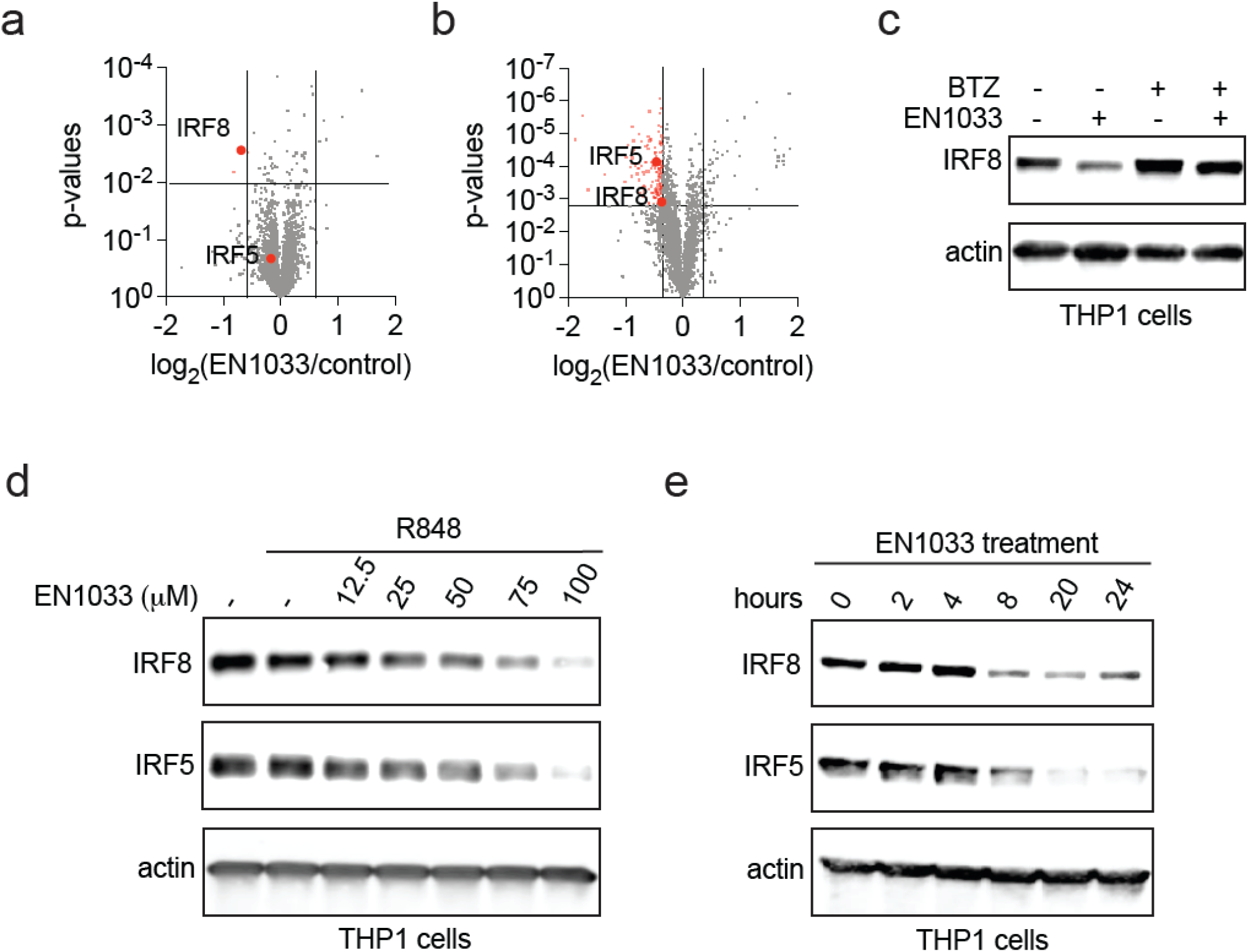
Validation of EN1033 as an IRF5 Degrader and Discovery of Effects on IRF8n. **(a, b)** Quantitative tandem mass tagging (TMT)-based proteomic profiling of EN1033 treatment. THP1 cells were treated with DMSO vehicle or EN1033 (100 μM) for 15 h **(a)** or 24 h **(b)**. IRF5 and IRF8 are highlighted. **(c)** Proteasome-dependence of IRF8 degradation. THP1 cells were pre-treated with DMSO vehicle or BTZ (500 nM) 1 h prior to treatment with DMSO vehicle or EN1033 (100 μM) for 18 h, after which IRF8 and loading control actin levels were assessed by SDS/PAGE and Western blotting. **(d)** Dose-response of IRF5 and IRF8 degradation. THP1 cells were stimulated with DMSO or R848 (10 µM) for 1 h prior to treatment of cells with DMSO vehicle or EN1033 for 24 h, after which IRF8, IRF5, and loading control actin levels were assessed by SDS/PAGE and Western blotting. **(e)** Time-course of IRF5 and IRF8 degradation. THP1 cells were treated with EN1033 (100 μM) and IRF8, IRF5, and loading control actin levels were assessed by SDS/PAGE and Western blotting. Data in **(a,b)** are from n=3 biologically independent replicates per group and proteomic data can be found in **Table S2**. Blots in (**c,d,e)** are representative of n=3 biologically independent replicates per group.

### Further Characterization of EN1033 with an Alkyne Functionalized Probe and Chemoproteomic Profiling

To further characterize EN1033 and its interactions with IRF8 and IRF5, we synthesized an alkyne-functionalized “clickable” probe analog of EN1033, TH3-189 **(Figure 3a)**. TH3-189 showed comparable degradation activity against IRF8 and IRF5 in THP1 cells **(Figure 3b)**. We next demonstrated that TH3-189 covalently labeled purified human IRF8 and IRF5 protein *in vitro* in a dose-responsive manner, as assessed by visualizing protein labeling by copper-catalyzed azide-alkyne cycloaddition (CuAAC) conjugation of an azide-functionalized rhodamine fluorophore on an SDS/PAGE gel **(Figure 3c-3d)**. With MS/MS analysis, we determined that EN1033 labeled both C223 and C385 on pure IRF8 protein **(Figure S5)**.

**Figure 3.**
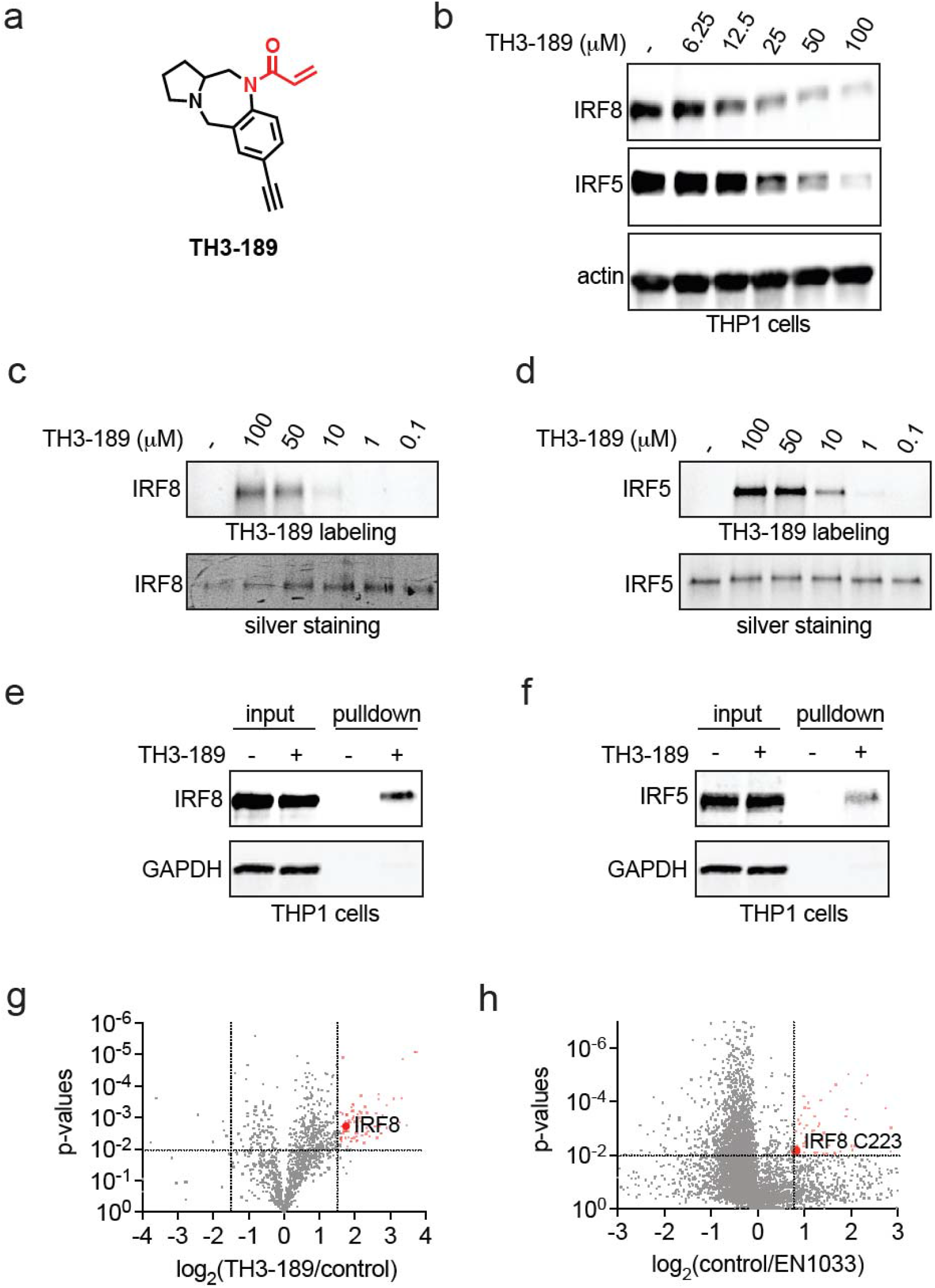
Further Characterization of EN1033 with an Alkyne Functionalized Probe and Through Chemoproteomic Profiling. **(a)** Structure of alkyne functionalized probe TH3-189 with the acrylamide warhead highlighted in red. **(b)** Dose-response of IRF5 and IRF8 loss. THP1 cells were treated with DMSO vehicle or EN1033 for 24 h after which IRF8, IRF5, and loading control actin levels were assessed by SDS/PAGE and Western blotting. **(c,d)** IRF8 and IRF5 pure protein labeling by TH3-189. Pure IRF8 **(c)** or IRF5 **(d)** protein was labeled with DMSO vehicle or TH3-189 (100 μM) for 1 h, after which an azide-functionalized rhodamine handle was appended onto probe-labeled proteins by CuAAC. Proteins were resolved by SDS/PAGE, and TH3-189 labeling was assessed by in-gel fluorescence, and loading was assessed by silver staining. **(e,f)** IRF8 and IRF5 pulldown with TH3-189 probe in cells. THP1 cells were treated with DMSO vehicle or TH3-189 (100 μM) for 4 h, after which probe-modified proteins from resulting lysates were appended to an azide-functionalized biotin enrichment handle by CuAAC, avidin-enriched, eluted, and IRF8, IRF5, and loading control actin levels from input and pulldown eluate were assessed by SDS/PAGE and Western blotting. **(g)** Chemoproteomic profiling of TH3-189 targets. THP1 cells were treated with DMSO vehicle or TH3-189 (100 μM) for 4 h, after which probe-modified proteins from resulting lysates were appended to an azide-functionalized biotin enrichment handle by CuAAC, avidin-enriched, eluted, tryptically digested, and analyzed by LC-MS/MS. Highlighted in red are proteins that were significantly enriched compared to the DMSO vehicle, and IRF8 is highlighted. **(h)** isoDTB-ABPP analysis of EN1033. THP1 cells were treated with DMSO vehicle or EN1033 (100 μM) for 6 h, after which the resulting lysates were labeled with an alkyne-functionalized iodoacetamide probe (IA-alkyne) (200 μM) for 1 h. Probe-labeled proteins were then appended to an isotopically light or heavy azide-functionalized desthiobiotin handle, enriched by streptavidin beads, eluted, tryptically digested, and analyzed by LC-MS/MS. Shown in red are cysteines in proteins that were significantly engaged with C223 or IRF8, highlighted in red. Blots and gels shown in **(b-f)** are representative of n=3 biologically independent replicates per group. Data in **(g,h)** are from n=3 biologically independent replicates per group and chemoproteomic data can be found in **Table S3** and **Table S4**, respectively.

We next sought to assess whether EN1033 directly engages IRF8 and IRF5 in THP1 cells. We treated cells with TH3-189, followed by CuAAC-mediated conjugation of an azide-functionalized biotin enrichment handle to the resulting lysates, and performed a pulldown of probe-labeled proteins. The results demonstrated significant enrichment of both IRF8 and IRF5 in THP1 cells **(Figure 3e-3f)**.

To further assess the proteome-wide selectivity of EN1033, we performed two orthogonal chemoproteomic profiling experiments. We first evaluated proteins significantly enriched by our TH3-189 probe. IRF8 was enriched considerably in TH3-189-treated cells compared to vehicle-treated controls, along with 81 other proteins that were also enriched among the >1400 proteins quantified **(Figure 3g; Table S3)**. Subsequent isotopic desthiobiotin-ABPP (isoDTB-ABPP) cysteine chemoproteomic profiling revealed significant engagement of C223 on IRF8 with 60 other off-targets **(Figure 3h; Table S4)** ^13,14,27^. In both chemoproteomic experiments, we did not detect IRF5 or identify IRF5 engagement, possibly because EN1033 or TH3-189 engaged IRF8 less than IRF8, or because indirect effects, perhaps downstream of IRF8, were affecting IRF5 levels. Nonetheless, our chemoproteomic profiling data further indicated that EN1033 more robustly engages IRF8 than IRF5 and that EN1033 or its analog TH3-189 is moderately selective across the proteome.

### Assessing Mechanism of IRF5 and IRF8 Degradation by EN1033

Given that our data thus far have demonstrated that EN1033 or its analog TH3-189 engages IRF5 and IRF8 *in vitro* and in cells, and degrades both proteins, we next sought to further assess target engagement and the mechanism of degradation. Cellular thermal shift assay (CETSA)^28^ was performed to determine the thermal stability of IRF8 and IRF5 upon EN1033 treatment. We observed significant destabilization of both IRF8 and IRF5, but not of the unrelated control protein actin, in THP1 cells treated with EN1033. The CETSA revealed a more substantial destabilization of IRF8 compared to IRF5, further supporting our chemoproteomic and proteomic findings, which indicate more acute and pronounced effects on IRF8 **(Figure 4a)**. These data also suggest that EN1033 may act similarly to our previously identified destabilizing degraders against MYC, CTNNB1, and AR-V7, through covalent binding and direct thermodynamic destabilization of IRF5 and IRF8 ^21-23^.

**Figure 4.**
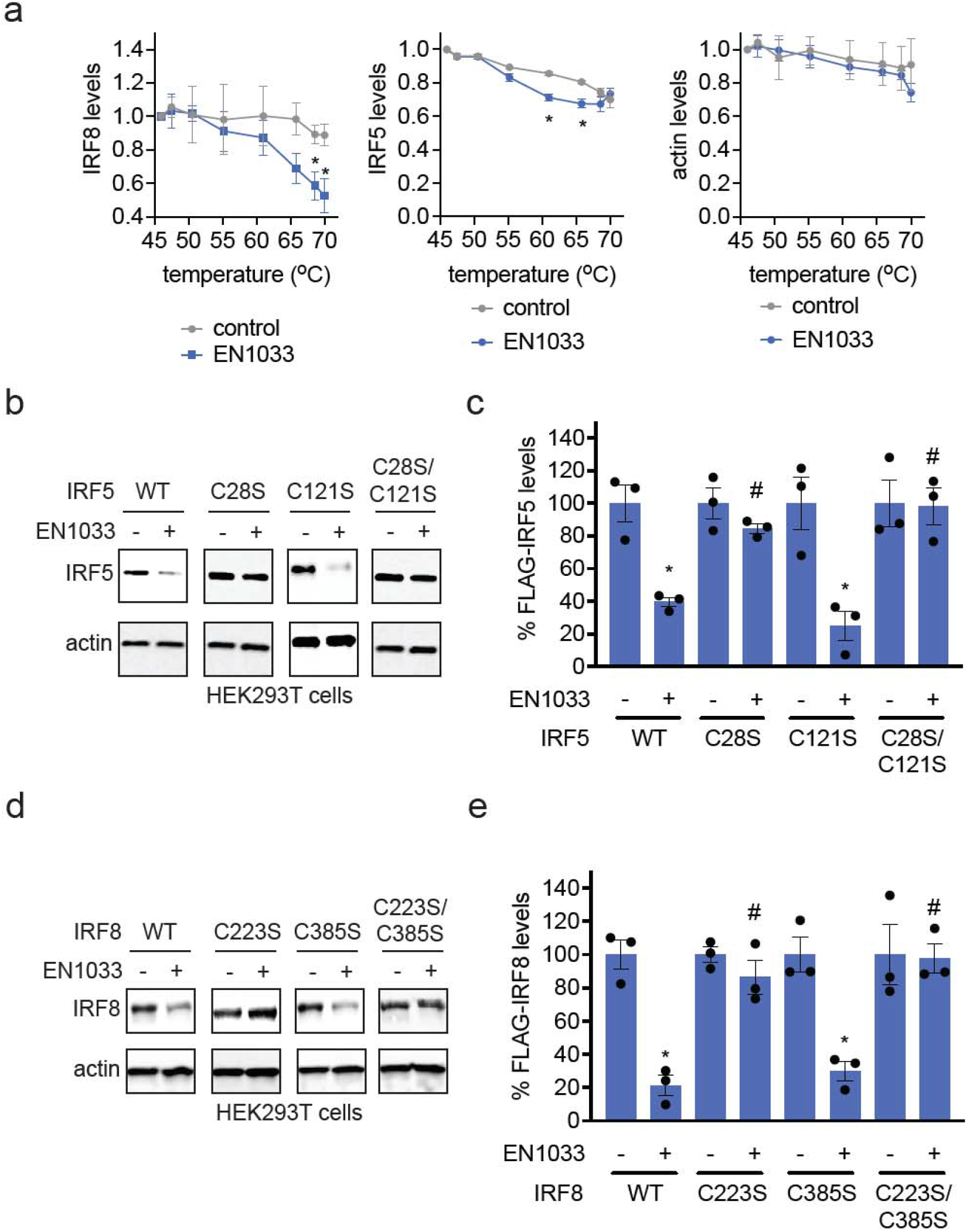
Assessing Mechanism of IRF5 and IRF8 Degradation by EN1033. **(a)** CETSA analysis of EN1033. THP1 cells were treated with DMSO vehicle or EN1033 (100 μM) for 4 h, after which cell lysates were heated to the designated temperatures, insoluble proteins were precipitated, and IRF8, IRF5, and control protein actin levels were assessed by Western blotting and quantified by densitometry. **(b,c,d,e)** IRF5 or IRF8 degradation by EN1033 in FLAG-IRF5 or FLAG-IRF8 WT or mutant expressing cells. HEK293T cells expressing FLAG-IRF5 WT, C28S, C121S, or C28S/C121S mutant were treated with DMSO vehicle or EN1033 (50 μM) for 24 h **(b,c)**, or FLAG-IRF8 WT, C223S, C385S, or C223S/C385S mutant **(d,e)** were treated with DMSO vehicle or EN1033 (50 μM) for 18 h, after which FLAG-IRF5, FLAG-IRF8, or loading control actin levels were assessed by SDS/PAGE and Western blotting and quantified in **(c,e)**. Data in **(a-e)** are from n=3 biologically independent replicates per group. Graphs in **(a)** show average ± sem. Bar graphs in **(c,e)** show individual replicate values and average ± sem. Significance in **(a,c,e)** is shown as *p<0.05 compared to vehicle-treated controls for each group and #p<0.05 compared to EN1033-treated FLAG-IRF5 WT or FLAG-IRF8 WT-expressing cells.

Considering the numerous off-target effects of EN1033, we sought to confirm that the degradation of IRF5 and IRF8 was mediated by direct engagement of cysteines on these proteins, rather than by indirect mechanisms arising from off-target action. Since EN1033 binds to C28 and C121 on IRF5 and to C223 and C385 on IRF8 *in vitro*, we mutated both respective cysteines to serines to assess which sites may be responsible for the destabilization of IRF5 and IRF8 **(Figure 4b-4e)**. With IRF5, we observed complete rescue of IRF5 degradation in C28S and C28S/C121S mutant IRF5-expressing cells, but not in C121S-expressing cells, indicating that C28 is likely the primary site for EN1033-mediated degradation of IRF5 **(Figure 4b-4c)**. Correspondingly, with IRF8, we observed complete rescue of IRF8 loss in the C223S and C223S/C385S mutant, but not in the C385S mutant IRF8-expressing cells, indicating that C223 is the primary site for EN1033-mediated degradation of IRF8 **(Figure 4d-4e)**. Our data thus demonstrate that EN1033 degrades IRF5 and IRF8 by targeting C28 and C223, respectively.

Nonetheless, taken together, our data suggest that EN1033 engages and degrades IRF8 more robustly than IRF5. As such, we sought to further understand if there was any mechanistic connection between IRF8 and IRF5 abundance. We demonstrated that EN1033 continues to degrade IRF8 in IRF5 KO cells stimulated with R848, suggesting that IRF5 does not likely regulate IRF8 **(Figure S6a)**. In contrast, we observed that IRF8 knockdown led to a marked reduction in IRF5 levels, indicating that IRF8 can regulate IRF5 in THP1 cells **(Figure S6b)**. These results are consistent with previous reports of IRF8 regulation of IRF5^29^. Thus, while we demonstrate that the EN1033-mediated IRF5 loss can be rescued, at least at acute 24 h timepoints, upon mutagenesis of C28, indicating direct action of our molecule, the IRF5 loss we observe could also be occurring through primary engagement and degradation of IRF8, leading to downstream transcriptional effects upon IRF5, particularly at later timepoints.

Further demonstrating functional IRF5/8 loss, we showed that R848-stimulated cell-surface levels of an IRF5/8-mediated marker of inflammation and monocyte lineage specification, CD83, were significantly downregulated upon EN1033 treatment, alongside IRF5 levels, using immunofluorescence approaches **(Figure S7a-S7b)**.

### Transcriptomic Profiling of EN1033

We next performed RNA sequencing on EN1033-treated THP1 cells to assess whether we observed global transcriptional changes consistent with inhibiting IRF5 and IRF8 function **(Figure 5a-5b; Table S5)**. We observed 809 and 448 transcripts significantly downregulated and upregulated by >2-fold, respectively **(Figure 5a; Table S5)**. Pathway analysis of these differentially expressed transcripts revealed several key inflammatory signaling nodes regulated by IRF5 and IRF8 that were significantly enriched, including IL-10, IL-4, IL-13, IFN-γ, cytokine, interleukin, chemokine, and IL-18 signaling **(Figure 5b)**. The transcriptomic changes elicited by EN1033 treatment revealed distinct yet overlapping immune signatures dependent on IRF5 and IRF8. IRF5-specific downregulation was notably evident in inflammatory cytokines and chemokines, including IL1A, IL1B, IL6, IL23A, and CCL2, reflecting IRF5’s central role in macrophage-driven inflammatory signaling, Th17 differentiation, and innate immune responses ^6,30^ **(Figure 5a, Table S5)**. Conversely, we observed downregulated genes that were explicitly controlled by IRF8, including its role in antigen presentation, dendritic cell activation, THP1-polarizing cytokine production, and lymphocyte trafficking, such as IRF8 itself, CD1C, CD83, CIITA, TLR8, CCR7, and the cytokines IL12B, CXCL9, and CXCL14^31,32^. Importantly, we also observed gene signatures that could be regulated by both IRF5 and IRF8, including critical immune regulatory mediators and chemokines such as CXCL10, CXCR3, IL10, SOCS1, SOCS3, IL18R1, and STAT4 ^6,31^. Collectively, the overall gene signature is consistent with inhibition of IRF5 and IRF8-driven pro-inflammatory programs.

**Figure 5.**
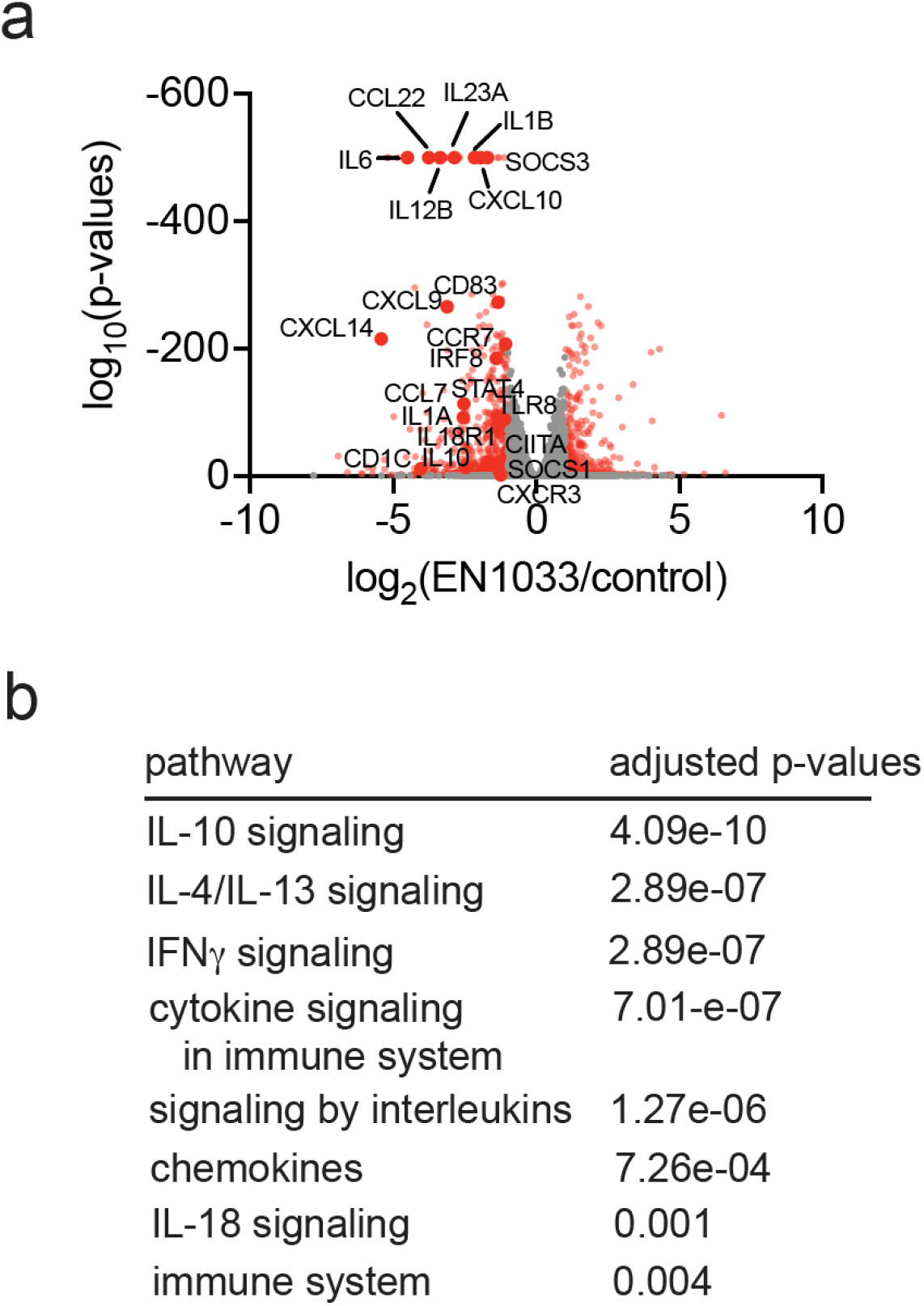
Transcriptomic profiling of EN1033. THP1 cells were stimulated with R848 (10 μM) and treated with DMSO vehicle or EN1033 (50 μM) for 24 h, after which RNA from cells was subjected to RNA sequencing and quantification. Shown in **(a)** are significantly altered transcripts with EN1033 treatment, with representative IRF5 and IRF8 target genes designated with gene names. Pathway analysis from significantly downregulated genes is shown in **(b)**. Full data are presented in **Table S5**. Data are from n=3 biologically independent replicates per group. P-values less than 1e-500 were capped at 1e-500.

### Improving Potency of EN1033

Given the poor potency of EN1033 against both IRF5 and IRF8, we sought to explore structure-activity relationships of our hit scaffold. We profiled 21 analogs of EN1033 for their ability to degrade IRF5 in a HiBiT-IRF5-expressing THP1 cell line **(Figure 6a-6b, Figure S8)**. We found that analog TH-B10 was significantly more potent at degrading IRF5, with a DC_50_ and D_max_ of 5.9 μM and 99.9 %, respectively, compared to EN1033, with a DC_50_ and D_max_ of 52 μM and 67.6 %, and other analogs **(Figure 6a-6b, Figure S8, Figure 7a)**. By Western blotting, measuring loss of endogenous IRF8 and IRF5, TH-B10 treatment led to a dose-responsive loss of both IRF8 and IRF5, with DC_50_ values of 2.2 and 6.4 μM, respectively **(Figure 7b-7c)**. Time-course studies clearly showed that TH-B10 reduced IRF8 levels more rapidly than IRF5 in cells **(Figure 7d-7e)**. Unlike in EN1033, we showed that IRF8 loss was proteasome-dependent, whereas IRF5 loss was not, indicating that TH-B10 may act by targeting IRF8 and that IRF5 downregulation was secondary through transcriptional downregulation **(Figure S9a-S9b)**. At earlier treatment times of 12 h, we observed robust loss of IRF8 over IRF5 with TH-B10 treatment, compared to EN1033 **(Figure S9c-S9d)**, and this loss of IRF8 was also observed in IRF5 knockout cells **(Figure S9e)**. Quantitative proteomic profiling showed a significant and selective reduction in IRF8 with TH-B10 treatment **(Figure 7f; Table S6)**. We also demonstrated dose-responsive inhibition of IRF5 transcriptional reporter activity in a R848- and TLR7/8-dependent manner, which was attenuated in IRF5 knockout cells **(Figure 7g)**. We also showed that TH-B10 only modestly inhibited IFN-α-mediated transcriptional reporter activity, not in an IRF5-dependent manner **(Figure 7h)**.

**Figure 6.**
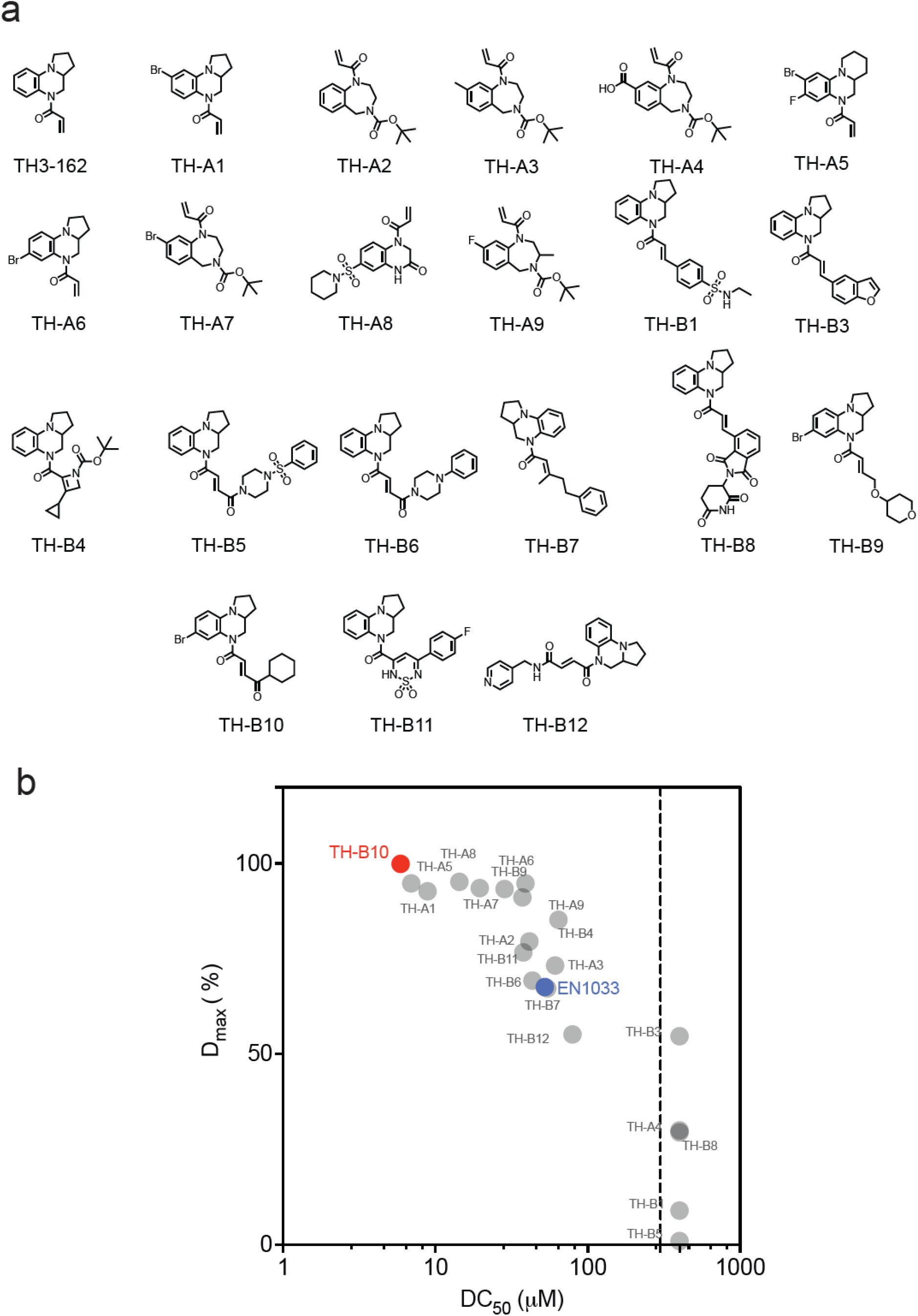
Structure-activity relationship of EN1033. a) Analogs of EN1033. b) HiBiT-IRF5 THP1 cells were pretreated with DMSO vehicle or compound in dose-response experiments for 24 h, and HiBiT-IRF5 levels were detected by luminescence. D_max_ and EC_50_ of degradation were calculated and plotted. Data are from n = 3 biologically independent replicates/group.

**Figure 7.**
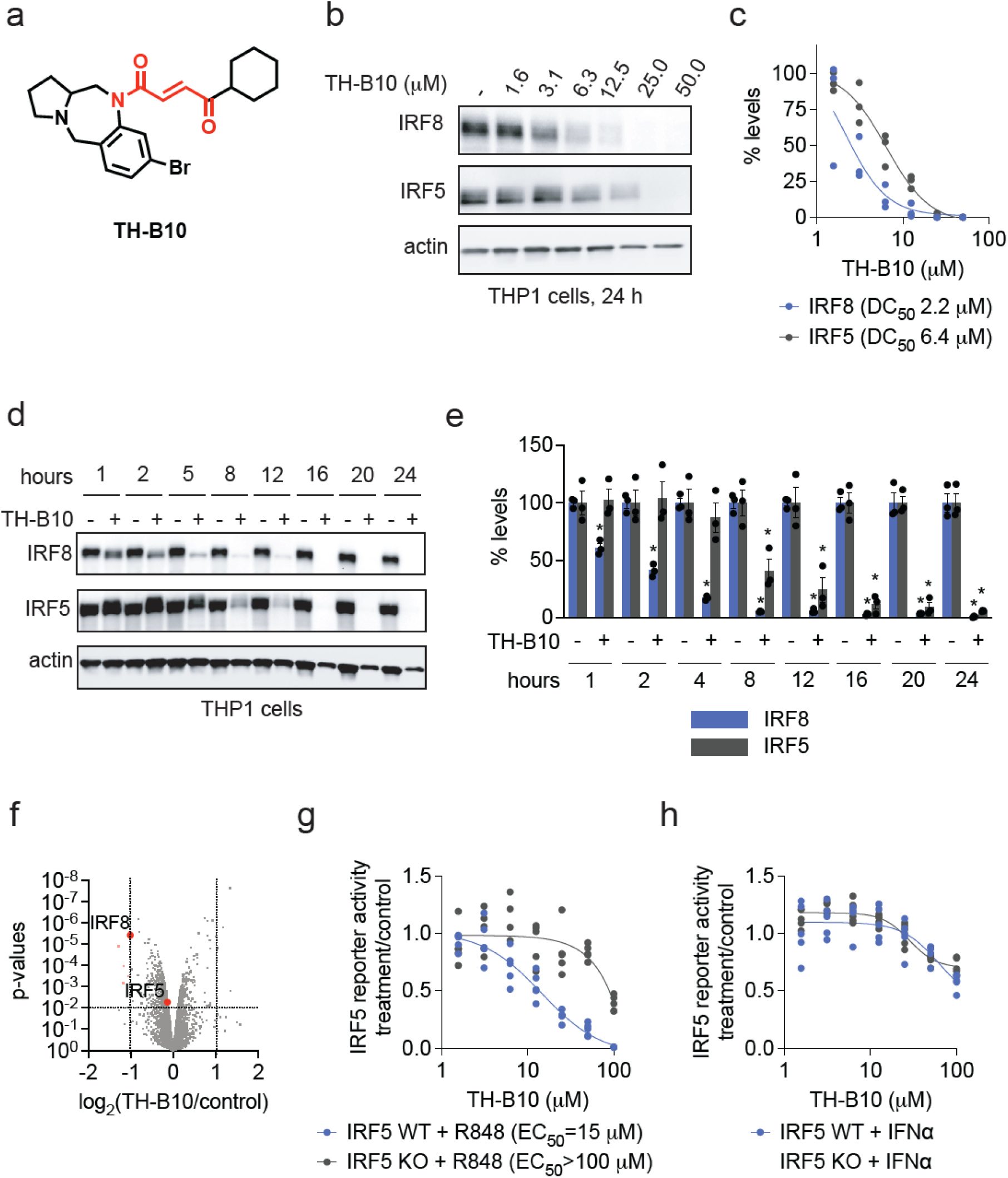
Characterization of optimized compound TH-B10. **(a)** Structure of TH-B10. **(b,c)** Dose-response of IRF8 and IRF5 loss. THP1 cells were treated with DMSO vehicle or TH-B10 for 24 h, after which IRF8, IRF5, and loading control actin levels were assessed by Western blotting **(b)** and quantified **(c). (d,e)** Time-course of TH-B10-mediated IRF8 and IRF5 degradation. THP1 cells were treated with a DMSO vehicle or TH-B10 (50 µM) at designated times, and IRF8, IRF5, and loading control actin levels were assessed by Western blotting and quantified **(e). (f)** TMT-based proteomic profiling of TH-B10 treatment. THP1 cells were treated with DMSO vehicle or TH-B10 (10 μM) for 12 h. IRF8 and IRF5 are highlighted. **(g,h)** IRF5 luciferase reporter activity. IRF5 WT or KO THP1 cells expressing an IRF5 luciferase reporter were stimulated with R848 (10 µM) **(g)** or IFNα (1000 U/mL) **(h)** and treated with DMSO vehicle or TH-B10 for 12 h, after which luciferase activity was assessed. Blots and gels in **(b,d)** are representative of n=3 biologically independent replicates per group. Graphs in **(e,g,h)** show individual replicate values and in **(e)** average ± sem. Significance expressed as *p<0.05 compared to vehicle-treated controls in **(e)**. Data in **(f)** are from n=3 biologically independent replicates per group and proteomic data can be found in **Table S6**. Data in **(g,h)** are from n=3-4 biologically independent replicates per group.

Given that the IRF5 loss was not proteasome-dependent degradation, we focused on the mechanistic validation of TH-B10-mediated IRF8 degradation. As observed with EN1033, TH-B10-mediated IRF8 degradation is attenuated in IRF8 C223S-expressing cells and is not further rescued by the double C223S/C385S mutant **(Figure 8a-8b)**. Further supporting on-target activity, we showed that cell-surface CD83 was also significantly downregulated by TH-B10 treatment in THP1 cells **(Figure S10a-S10b)**. Transcriptomic profiling of TH-B10-treated cells also showed significant downregulation of expected genes involved in IRF5-mediated inflammatory responses, as well as in TNF and IFN responses **(Figure 8c-8d; Figure S11; Table S7)**.

**Figure 8.**
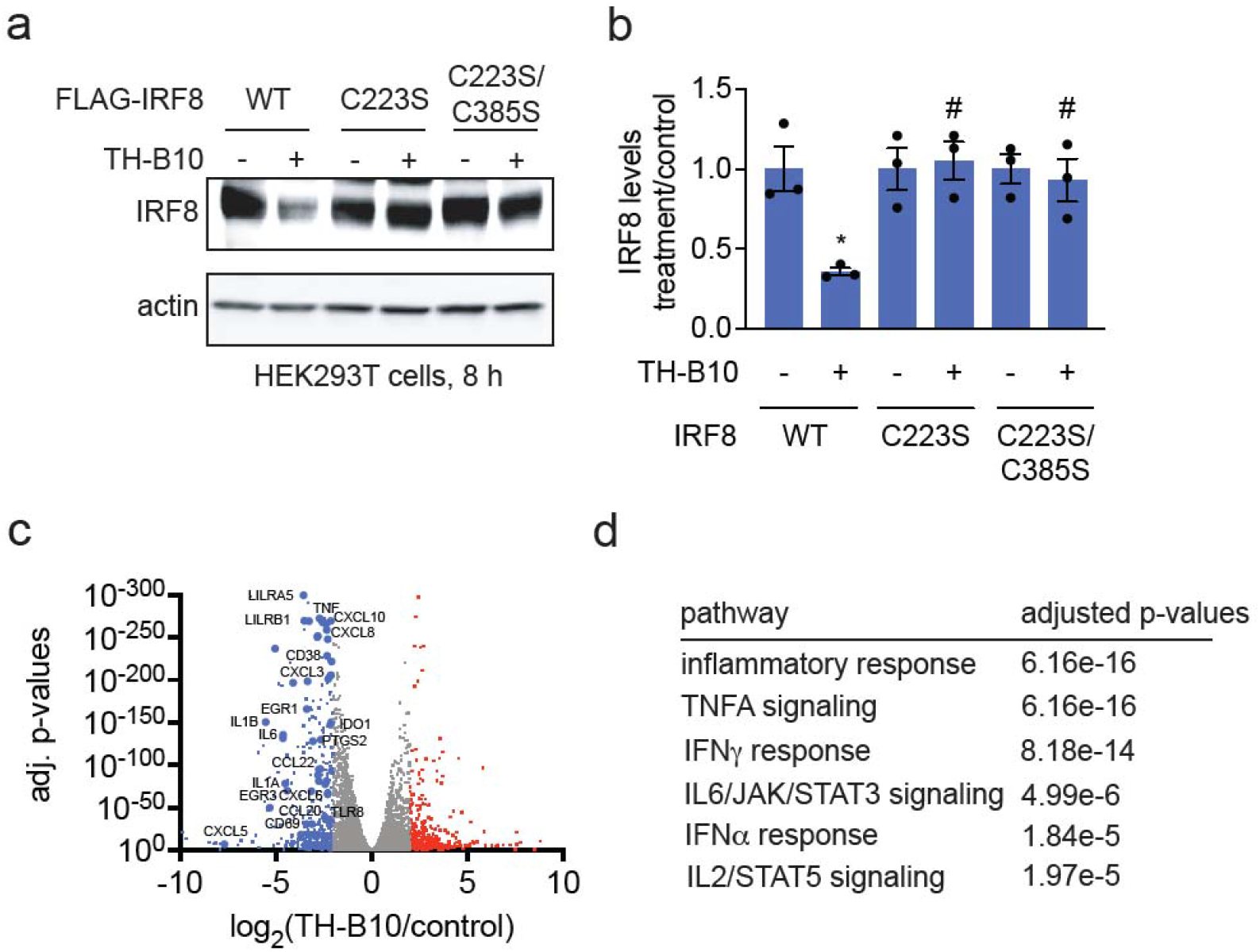
Degradation of IRF8 and inflammatory immune response by TH-B10. **(a,b)** IRF8 degradation by TH-B10 in FLAG-IRF8 WT or mutant expressing cells. HEK293T cells expressing FLAG-IRF8 WT, C223S, C385S, mutants were treated with DMSO vehicle or TH-B10 (10 μM) for 8 h, after which FLAG-IRF8 or loading control actin levels were assessed by SDS/PAGE and Western blotting **(a)** and quantified in **(b). (c,d)** Transcriptomic profiling of TH-B10. THP1 cells were stimulated with R848 (10 μM) and treated with DMSO vehicle or TH-B10 (10 μM) for 12 h, after which RNA from cells was subjected to RNA sequencing and quantification. Shown in **(c)** are significantly altered transcripts with TH-B10 treatment, with representative IRF5 and IRF8 target genes designated with gene names. Pathway analysis from significantly downregulated genes is shown in **(d)**. Data in **(a)** is representative of n=3 biologically independent replicates. Bar graphs in **(b)** show individual replicate values and average ± sem. Significance in **(b)** is shown as *p<0.05 compared to vehicle-treated controls for each group and #p<0.05 compared to TH-B10-treated FLAG-IRF8 WT-expressing cells. Full data from **(c,d)** are presented in **Table S7**. Data from **(c)** are from n=3 biologically independent replicates per group.

Overall, we have generated TH-B10, a more potent molecule that now covalently targets IRF8, leading to subsequent downregulation of IRF5, inhibition of inflammatory transcriptional activity, and reduced expression of inflammatory target genes.

## Discussion

In this study, we report the discovery of a covalent, monovalent degrader, EN1033, that induces proteasome-dependent degradation of the immune regulatory transcription factors IRF5 and IRF8. Using a combination of chemical proteomics, cysteine-reactive probe screening, transcriptional reporter assays, mutational validation, and cellular engagement studies, we show that EN1033 engages and destabilizes IRF5 and IRF8 through covalent modification of specific cysteine residues—C28 on IRF5 and C223 on IRF8— leading to loss of protein levels and inhibition of inflammatory transcriptional activity. Unexpectedly, EN1033 exhibited more potent and rapid engagement of IRF8 than IRF5, with functional consequences on IRF5 expression downstream of IRF8 degradation, revealing a previously underappreciated regulatory axis between these two key transcriptional drivers of inflammation. Further structure-activity relationships yielded an optimized compound, TH-B10, that targets IRF8 more selectively and robustly, and downregulates IRF5, thereby inhibiting inflammatory transcriptional programming.

These findings represent one of the first demonstrations of a direct-acting, covalent degrader against IRF8—a transcription factor historically considered undruggable due to its intrinsic disorder and lack of defined binding pockets. More broadly, this work expands the utility of covalent chemoproteomic and ABPP strategies in targeting structurally disordered immune transcription factors. Prior studies have demonstrated the feasibility of this approach against oncogenic transcription factors, such as MYC, β-catenin, AR-V7, and FOXA1^16,17,21–23^. This study uniquely extends it into the immunological space, targeting IRFs that are deeply embedded in innate immune signaling, macrophage activation, and the pathophysiology of autoimmune diseases. The dual degradation of IRF5 and IRF8 by EN1033 and the degradation of IRF8 by TH-B10 not only highlights its pharmacological utility but also provides a chemical probe to dissect transcriptional circuitry in innate immune pathways.

Nonetheless, several caveats warrant consideration. First, while the degrader showed proteasome-dependent degradation and cysteine-specific engagement in both in vitro and cellular settings, global chemoproteomic profiling revealed off-target activity, with dozens of additional protein interactions, raising questions about its selectivity in complex proteomes. Second, although mutagenesis of target cysteines confirmed the degrader’s direct mechanism of action, our data also suggest that IRF5 downregulation may also occur indirectly through IRF8 loss. These downstream effects complicate the distinction between primary degradation and secondary downregulation events and will require further temporal and mechanistic resolution. Third, given the intrinsic reactivity of electrophilic acrylamides, further medicinal chemistry optimization will be necessary to enhance selectivity, reduce off-target engagement, and optimize potency.

Finally, functional in vivo validation and exploration of therapeutic efficacy in models of inflammation or autoimmunity remain to be established.

Overall, this study establishes EN1033 and TH-B10 as early-stage chemical tools and pathfinder molecules^33^ for modulating IRF8 and IRF5, and more broadly underscores the promise of covalent destabilizing degraders in tackling transcription factors that have long been considered intractable to traditional small-molecule approaches. These findings open new avenues for the development of precision therapeutics in immune-driven diseases by leveraging the covalent ligandability of transcriptional regulators.

## Supporting information

Tabls S5

## Acknowledgment

We thank the members of the Nomura Research Group and Octant Bio for their critical review of the manuscript. This work was also supported by Octant Bio, the National Science Foundation Molecular Foundations for Biotechnology (MFB) grant (2127788), the UC Berkeley Molecular Therapeutics Initiative (MTI), the Mark Foundation for Cancer Research ASPIRE Award, the Mark Foundation for Cancer Research Momentum Postdoctoral Fellowship, Bakar Fellows Award, and the National Institutes of Health (R35CA263814, R01CA240981). We also thank Hasan, Lund, and the UC Berkeley NMR facility in the College of Chemistry (CoC-NMR) for spectroscopic assistance. Instruments in the College of Chemistry NMR facility are partly supported by NIH S10OD024998.

## Associated Content

### Supporting Information

The Supporting Information is available free of charge.

Supporting Table Legends, Supporting Figures, Materials and Methods, Synthetic Methods and Characterization (PDF).

