## Supporting Information for "Covalent Modulators of Immune Regulatory Transcription Factors IRF8 and IRF5"

[1] Department of Chemistry. University of California, Berkeley, Berkeley, CA 94720 USA

[2] Department of Molecular and Cell Biology, University of California, Berkeley, Berkeley, CA 94720 USA

[3] Innovative Genomics Institute, Berkeley, CA 94720 USA

[4] Octant Bio, Emeryville, CA 94608 USA

**Supplemental Table Legends**

**Table S1. Covalent ligand screen in HiBiT-IRF5 expressing THP1 cells.** Shown are the structures and compound names, as well as screening data of all cysteine-reactive covalent ligands screened. HiBiT-IRF5 expressing THP1 cells were treated with DMSO vehicle or covalent ligand (50 μM) for 24 h, after which HiBiT-IRF5 was detected by luminescence through detection with LgBiT and normalized to DMSO control.

**Table S2. Quantitative tandem mass tagging (TMT)-based proteomic profiling of EN1033 treatment.** THP1 cells were treated with DMSO vehicle or EN1033 (100 μM) for 15 h **(Tab 1)** or 24 h **(Tab 2).**

**Table S3. Chemoproteomic profiling of TH3-189 targets.** THP1 cells were treated with DMSO vehicle or TH3-189 (100 μM) for 4 h, after which probe-modified proteins from resulting lysates were appended to an azide-functionalized biotin enrichment handle by CuAAC, avidin-enriched, eluted, tryptically digested, and analyzed by LC-MS/MS. Data are from n=3 biologically independent replicates per group.

**Table S4. isoDTB-ABPP analysis of EN1033.** THP1 cells were treated with DMSO vehicle or EN1033 (100 μM) for 6 h, after which the resulting lysates were labeled with an alkyne-functionalized iodoacetamide probe (IA-alkyne) (200 μM) for 1 h. Probe-labeled proteins were then appended to an isotopically light or heavy azide-functionalized desthiobiotin handle, enriched by streptavidin beads, eluted, tryptically digested, and analyzed by LC-MS/MS. Data are from n=3 biologically independent replicates per group.

**Table S5. Transcriptomic profiling of EN1033.** THP1 cells were stimulated with R848 (10 μM) and treated with DMSO vehicle or EN1033 (50 μM) for 24 h, after which RNA from cells was subjected to RNA sequencing and quantification. Data are from n=3 biologically independent replicates per group. Tab 1 shows differential expression results from comparison of DMSO versus treatment. Tab 2 has estimated counts adjusted for library size.

**Table S6. Quantitative tandem mass tagging (TMT)-based proteomic profiling of TH-B10 treatment.** THP1 cells were treated with DMSO vehicle or TH-B10 (10 μM) for 12 h.

**Table S7. Transcriptomic profiling of TH-B10.** THP1 cells were stimulated with R848 (10 μM) and treated with DMSO vehicle or TH-B10 (10 μM) for 12 h, after which RNA from cells was subjected to RNA sequencing and quantification. Data are from n=3 biologically independent replicates per group. Tab 1 shows differential expression results from comparison of DMSO versus treatment. Tab 2 has pathway enrichment analysis.

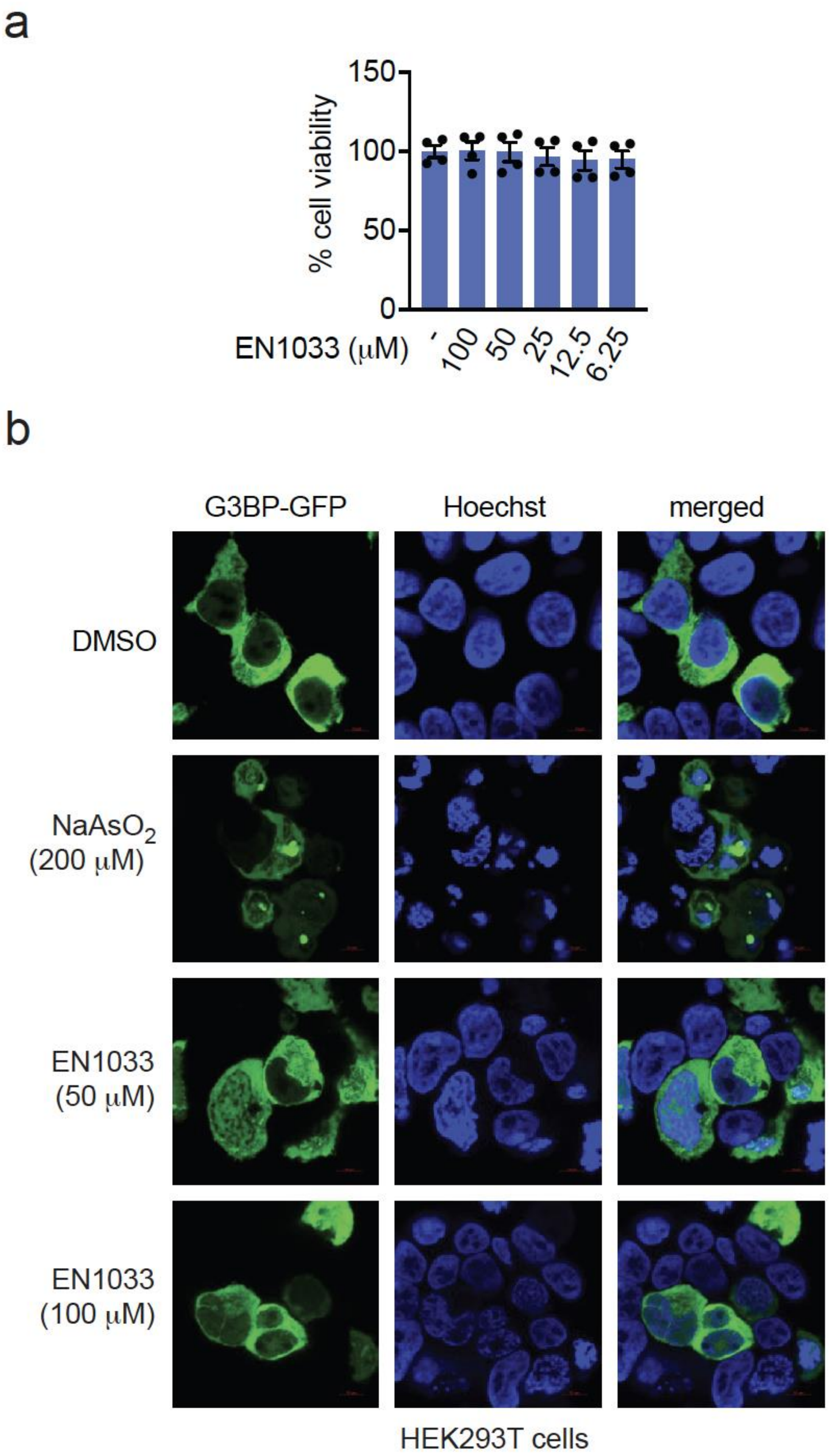


**Figure S1. Characterization of EN1033. (a)** THP1 cells were treated with DMSO vehicle or EN1033 for 24 h, and cell viability was assessed by CellTiter-Glo. **(b)** HEK293T cells were treated with DMSO vehicle, sodium arsenite, or EN1033 for 8 h, after which stress granule puncta were assessed by microscopy through read-out of G3BP-GFP. Hoechst was used to stain nuclei. Data in **(a)** show individual replicate values and average ± sem from n=4 biologically independent replicates per group. Data in **(b)** are representative of n=3 biologically independent replicates per group.

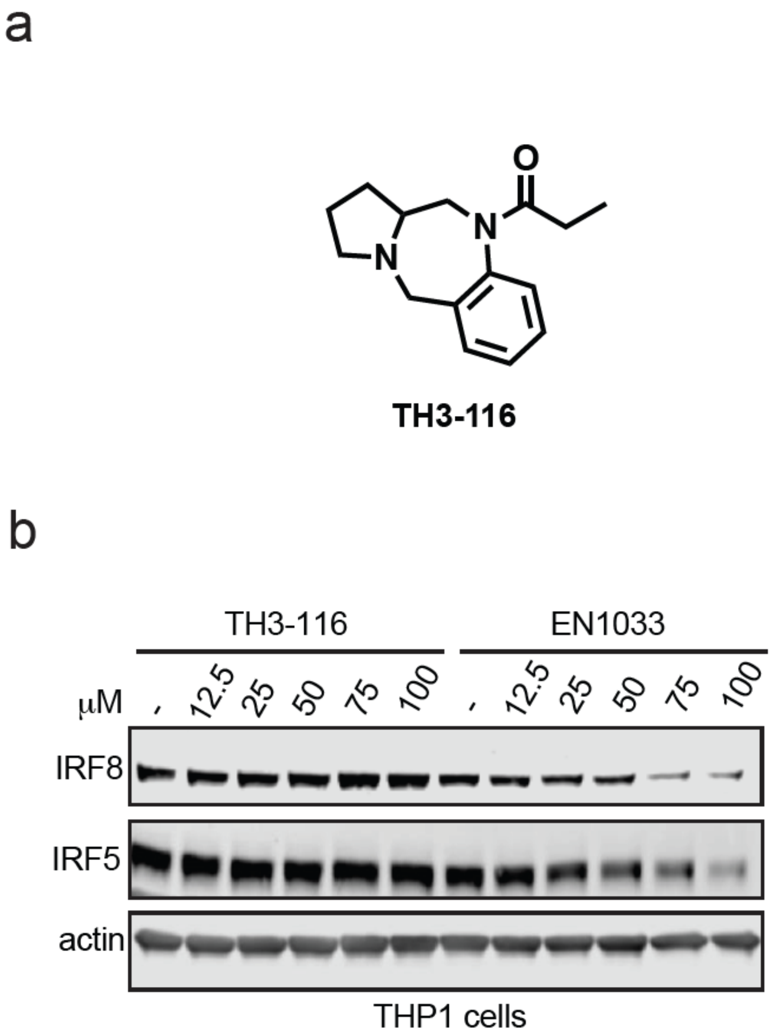


**Figure S2. Non-reactive control TH3-116 does not degrade IRF8 or IRF5. (a)** Structure of non-reactive analog TH3-116. **(b)** THP1 cells were treated with DMSO vehicle control, TH3-116, or EN1033 for 24 h after which IRF8, IRF5, and loading control actin levels were assessed by SDS/PAGE and Western blotting. Blot shown is representative of n=3 biologically independent replicates per group.

IRF5 C28

L K P W L V A Q V N S C* Q Y P G L Q W V N G E K

[M+H] 3000.5673, m/z 750.8973

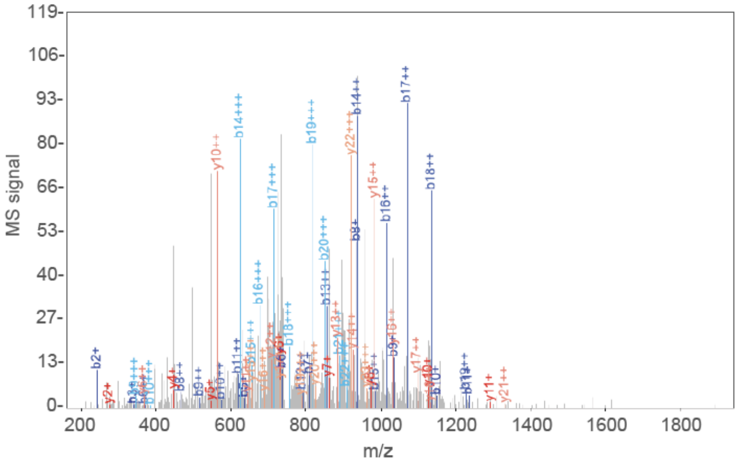


| b+ | b2+ | b3+ | # | Seq | # | y+ | y2+ | y3+ |
|---|---|---|---|---|---|---|---|---|
| 114.0913 | 57.5493 | 38.7020 | 1 | L | 24 | | | |
| 242.1863 | 121.5968 | 81.4003 | 2 | K | 23 | 2887.4832 | 1444.2452 | 963.1659 |
| 339.2391 | 170.1232 | 113.7512 | 3 | P | 22 | 2759.3883 | 1380.1978 | 920.4676 |
| 525.3184 | 263.1628 | 175.7776 | 4 | W | 21 | 2662.3355 | 1331.6714 | 888.1167 |
| 638.4024 | 319.7049 | 213.4723 | 5 | L | 20 | 2476.2562 | 1238.6317 | 826.0902 |
| 737.4709 | 369.2391 | 246.4951 | 6 | V | 19 | 2363.1721 | 1182.0897 | 788.3956 |
| 808.5080 | 404.7576 | 270.1742 | 7 | A | 18 | 2264.1037 | 1132.5555 | 755.3727 |
| 936.5665 | 468.7869 | 312.8604 | 8 | Q | 17 | 2193.0666 | 1097.0369 | 731.6937 |
| 1035.6350 | 518.3211 | 345.8832 | 9 | V | 16 | 2065.0080 | 1033.0076 | 689.0075 |
| 1149.6779 | 575.3426 | 383.8975 | 10 | N | 15 | 1965.9396 | 983.4734 | 655.9847 |
| 1236.7099 | 618.8586 | 412.9082 | 11 | S | 14 | 1851.8967 | 926.4520 | 617.9704 |
| 1582.8683 | 791.9378 | 528.2943 | 12 | C | 13 | 1764.8646 | 882.9360 | 588.9597 |
| 1710.9268 | 855.9671 | 570.9805 | 13 | Q | 12 | 1418.7063 | 709.8568 | 473.5736 |
| 1873.9902 | 937.4987 | 625.3349 | 14 | Y | 11 | 1290.6477 | 645.8275 | 430.8874 |
| 1971.0429 | 986.0251 | 657.6858 | 15 | P | 10 | 1127.5844 | 564.2958 | 376.5330 |
| 2028.0644 | 1014.5358 | 676.6930 | 16 | G | 9 | 1030.5316 | 515.7694 | 344.1821 |
| 2141.1485 | 1071.0779 | 714.3877 | 17 | L | 8 | 973.5102 | 487.2587 | 325.1749 |
| 2269.2070 | 1135.1072 | 757.0739 | 18 | Q | 7 | 860.4261 | 430.7167 | 287.4802 |
| 2455.2864 | 1228.1468 | 819.1003 | 19 | W | 6 | 732.3675 | 366.6874 | 244.7940 |
| 2554.3548 | 1277.6810 | 852.1231 | 20 | V | 5 | 546.2882 | 273.6477 | 182.7676 |
| 2668.3977 | 1334.7025 | 890.1374 | 21 | N | 4 | 447.2198 | 224.1135 | 149.7448 |
| 2725.4192 | 1363.2132 | 909.1446 | 22 | G | 3 | 333.1769 | 167.0921 | 111.7305 |
| 2854.4618 | 1427.7345 | 952.1588 | 23 | E | 2 | 276.1554 | 138.5813 | 92.7233 |
| | | | 24 | K | 1 | 147.1128 | 74.0600 | 49.7091 |

IRF5 C121

I Y E V C* S N G P A P T D S Q P P E D Y S F G A G E E E E E E E E E L Q R

[M+H] 4273.8422, m/z 1425.2856

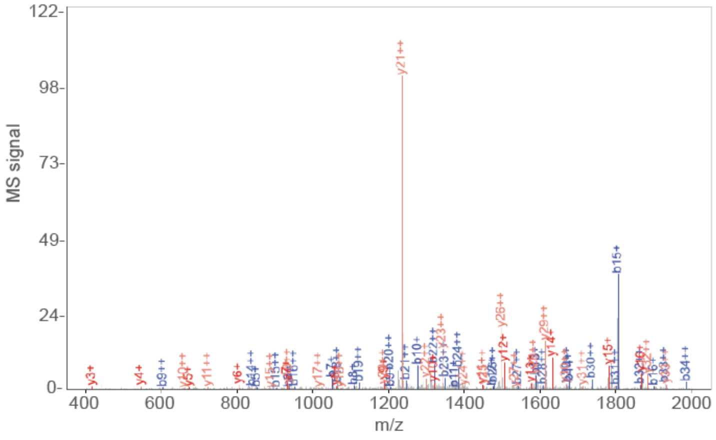


| b+ | b2+ | # | Seq | # | y+ | y2+ |
|---|---|---|---|---|---|---|
| 114.0913 | 57.5493 | 1 | I | 36 | | |
| 277.1547 | 139.0810 | 2 | Y | 35 | 4160.7582 | 2080.8827 |
| 406.1973 | 203.6023 | 3 | E | 34 | 3997.6949 | 1999.3511 |
| 505.2657 | 253.1365 | 4 | V | 33 | 3868.6523 | 1934.8298 |
| 851.4240 | 426.2156 | 5 | C | 32 | 3769.5838 | 1885.2956 |
| 938.4560 | 469.7317 | 6 | S | 31 | 3423.4255 | 1712.2164 |
| 1052.4990 | 526.7531 | 7 | N | 30 | 3336.3935 | 1668.7004 |
| 1109.5204 | 555.2639 | 8 | G | 29 | 3222.3505 | 1611.6789 |
| 1206.5732 | 603.7902 | 9 | P | 28 | 3165.3291 | 1583.1682 |
| 1277.6103 | 639.3088 | 10 | A | 27 | 3068.2763 | 1534.6418 |
| 1374.6631 | 687.8352 | 11 | P | 26 | 2997.2392 | 1499.1232 |
| 1475.7108 | 738.3590 | 12 | T | 25 | 2900.1864 | 1450.5969 |
| 1590.7377 | 795.8725 | 13 | D | 24 | 2799.1388 | 1400.0730 |
| 1677.7697 | 839.3885 | 14 | S | 23 | 2684.1118 | 1342.5595 |
| 1805.8283 | 903.4178 | 15 | Q | 22 | 2597.0798 | 1299.0435 |
| 1902.8811 | 951.9442 | 16 | P | 21 | 2469.0212 | 1235.0142 |
| 1999.9338 | 1000.4706 | 17 | P | 20 | 2371.9685 | 1186.4879 |
| 2128.9764 | 1064.9919 | 18 | E | 19 | 2274.9157 | 1137.9615 |
| 2244.0034 | 1122.5053 | 19 | D | 18 | 2145.8731 | 1073.4402 |
| 2407.0667 | 1204.0370 | 20 | Y | 17 | 2030.8462 | 1015.9267 |
| 2494.0987 | 1247.5530 | 21 | S | 16 | 1867.7828 | 934.3950 |
| 2641.1671 | 1321.0872 | 22 | F | 15 | 1780.7508 | 890.8790 |
| 2698.1886 | 1349.5979 | 23 | G | 14 | 1633.6824 | 817.3448 |
| 2769.2257 | 1385.1165 | 24 | A | 13 | 1576.6609 | 788.8341 |
| 2826.2472 | 1413.6272 | 25 | G | 12 | 1505.6238 | 753.3155 |
| 2955.2898 | 1478.1485 | 26 | E | 11 | 1448.6023 | 724.8048 |
| 3084.3324 | 1542.6698 | 27 | E | 10 | 1319.5597 | 660.2835 |
| 3213.3750 | 1607.1911 | 28 | E | 9 | 1190.5172 | 595.7622 |
| 3342.4176 | 1671.7124 | 29 | E | 8 | 1061.4746 | 531.2409 |
| 3471.4602 | 1736.2337 | 30 | E | 7 | 932.4320 | 466.7196 |
| 3600.5027 | 1800.7550 | 31 | E | 6 | 803.3894 | 402.1983 |
| 3729.5453 | 1865.2763 | 32 | E | 5 | 674.3468 | 337.6770 |
| 3858.5879 | 1929.7976 | 33 | E | 4 | 545.3042 | 273.1557 |
| 3971.6720 | 1986.3396 | 34 | L | 3 | 416.2616 | 208.6344 |
| 4099.7306 | 2050.3689 | 35 | Q | 2 | 303.1775 | 152.0924 |
| | | 36 | R | 1 | 175.1190 | 88.0631 |

**Figure S3. EN1033 site of modification on IRF5.** IRF5 pure protein (40 mg) was labeled with EN1033 (100 μM) for 1 h, followed by tryptic digestion of the protein and analysis of tryptic peptides by LC-MS/MS to identify EN1033-modified tryptic peptides. Shown are the MS/MS spectra of EN1033-modified tryptic peptides on C28 and C121 on the top and bottom, respectively.

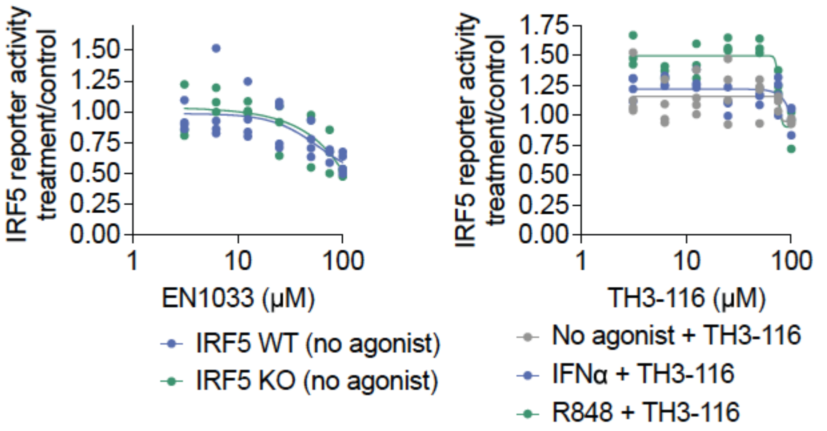


**Figure S4. IRF5 luciferase reporter activity.** IRF5 wildtype (WT) or knockout (KO) THP1 cells expressing an IRF5 luciferase reporter were either left unstimulated (no agonist) or cotreated with IFNa (1000 U/mL) **(g)** or R848 (10 μM) **(h)** and DMSO or vehicle EN1033 or TH3-116 for 24 h, after which luciferase activity was assessed. Shown are individual replicate data from n=3 biologically independent replicates.

IRF8 C223

L V G Q A T T T C* P E G C R

[M+H] 1711.8409, m/z 856.4241

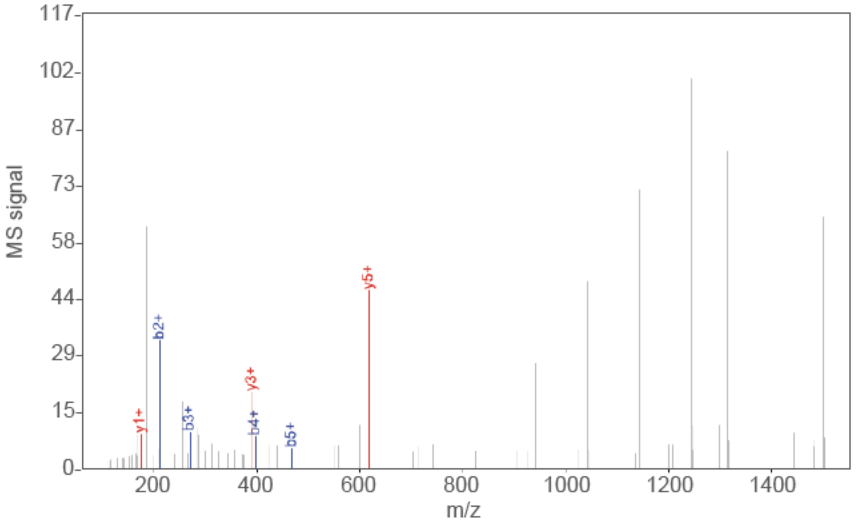


| b+ | # | Seq | # | y+ |
|---|---|---|---|---|
| 114.0913 | 1 | L | 14 | |
| 213.1598 | 2 | V | 13 | 1598.7569 |
| 270.1812 | 3 | G | 12 | 1499.6885 |
| 398.2398 | 4 | Q | 11 | 1442.6670 |
| 469.2769 | 5 | A | 10 | 1314.6084 |
| 570.3246 | 6 | T | 9 | 1243.5713 |
| 671.3723 | 7 | T | 8 | 1142.5236 |
| 772.4199 | 8 | T | 7 | 1041.4759 |
| 1094.5818 | 9 | C | 6 | 940.4283 |
| 1191.6346 | 10 | P | 5 | 618.2664 |
| 1320.6771 | 11 | E | 4 | 521.2137 |
| 1377.6986 | 12 | G | 3 | 392.1711 |
| 1537.7293 | 13 | C | 2 | 335.1496 |
| | 14 | R | 1 | 175.1190 |

IRF8 C385

S C* G A G S V M Q A P E E P P P D Q V F R

[M+H] 2478.1428, m/z 826.7191

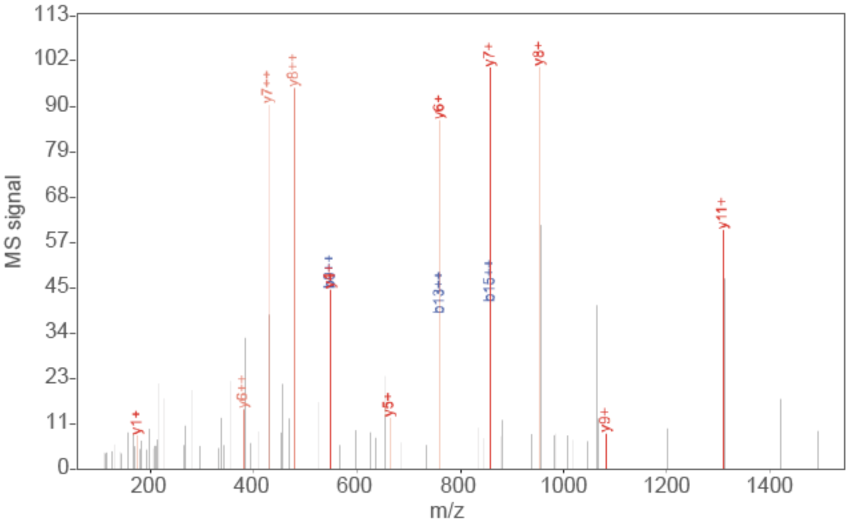


| b+ | b2+ | # | Seq | # | y+ | y2+ |
|---|---|---|---|---|---|---|
| 88.0393 | 44.5233 | 1 | S | 21 | | |
| 451.2009 | 226.1041 | 2 | C | 20 | 2391.1108 | 1196.0590 |
| 508.2224 | 254.6148 | 3 | G | 19 | 2027.9491 | 1014.4782 |
| 579.2595 | 290.1334 | 4 | A | 18 | 1970.9277 | 985.9675 |
| 636.2810 | 318.6441 | 5 | G | 17 | 1899.8905 | 950.4489 |
| 723.3130 | 362.1601 | 6 | S | 16 | 1842.8691 | 921.9382 |
| 822.3814 | 411.6944 | 7 | V | 15 | 1755.8370 | 878.4222 |
| 969.4168 | 485.2121 | 8 | M | 14 | 1656.7686 | 828.8880 |
| 1097.4754 | 549.2413 | 9 | Q | 13 | 1509.7332 | 755.3703 |
| 1168.5125 | 584.7599 | 10 | A | 12 | 1381.6747 | 691.3410 |
| 1265.5653 | 633.2863 | 11 | P | 11 | 1310.6375 | 655.8224 |
| 1394.6079 | 697.8076 | 12 | E | 10 | 1213.5848 | 607.2960 |
| 1523.6505 | 762.3289 | 13 | E | 9 | 1084.5422 | 542.7747 |
| 1620.7032 | 810.8553 | 14 | P | 8 | 955.4996 | 478.2534 |
| 1717.7560 | 859.3816 | 15 | P | 7 | 858.4468 | 429.7271 |
| 1814.8088 | 907.9080 | 16 | P | 6 | 761.3941 | 381.2007 |
| 1929.8357 | 965.4215 | 17 | D | 5 | 664.3413 | 332.6743 |
| 2057.8943 | 1029.4508 | 18 | Q | 4 | 549.3144 | 275.1608 |
| 2156.9627 | 1078.9850 | 19 | V | 3 | 421.2558 | 211.1315 |
| 2304.0311 | 1152.5192 | 20 | F | 2 | 322.1874 | 161.5973 |
| | | 21 | R | 1 | 175.1190 | 88.0631 |

**Figure S5. EN1033 site of modification on IRF8.** IRF8 pure protein (20 mg) was labeled with EN1033 (100 µM) for 1 h, followed by tryptic digestion of the protein and analysis of tryptic peptides by LC-MS/MS to identify EN1033-modified tryptic peptides. Shown are the MS/MS spectra of EN1033-modified tryptic peptides on C223 and C385 on the top and bottom, respectively.

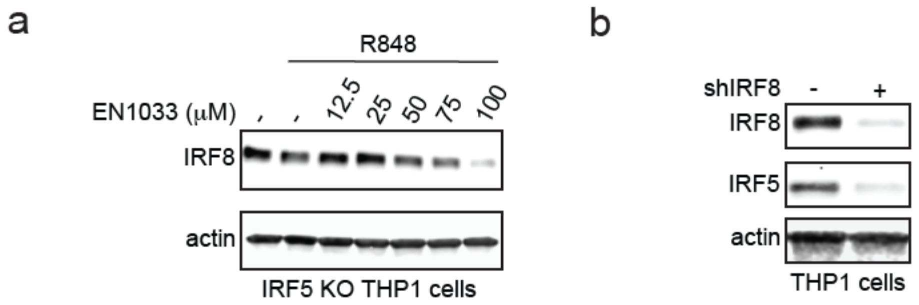


**Figure S6. Relationship between IRF8 and IRF5. (a)** IRF5 KO THP1 cells were treated with DMSO vehicle or stimulated with R848 (10 μM) for 1 h, and with EN1033 for 24 h. IRF8 and loading control actin levels were assessed by SDS/PAGE and Western blotting. **(b)** IRF8 and IRF5 and loading control actin levels were assessed by SDS/PAGE and Western blotting in stable shControl and shIRF8 THP1 cells. Blots shown are representative of n=3 biologically independent replicates per group.

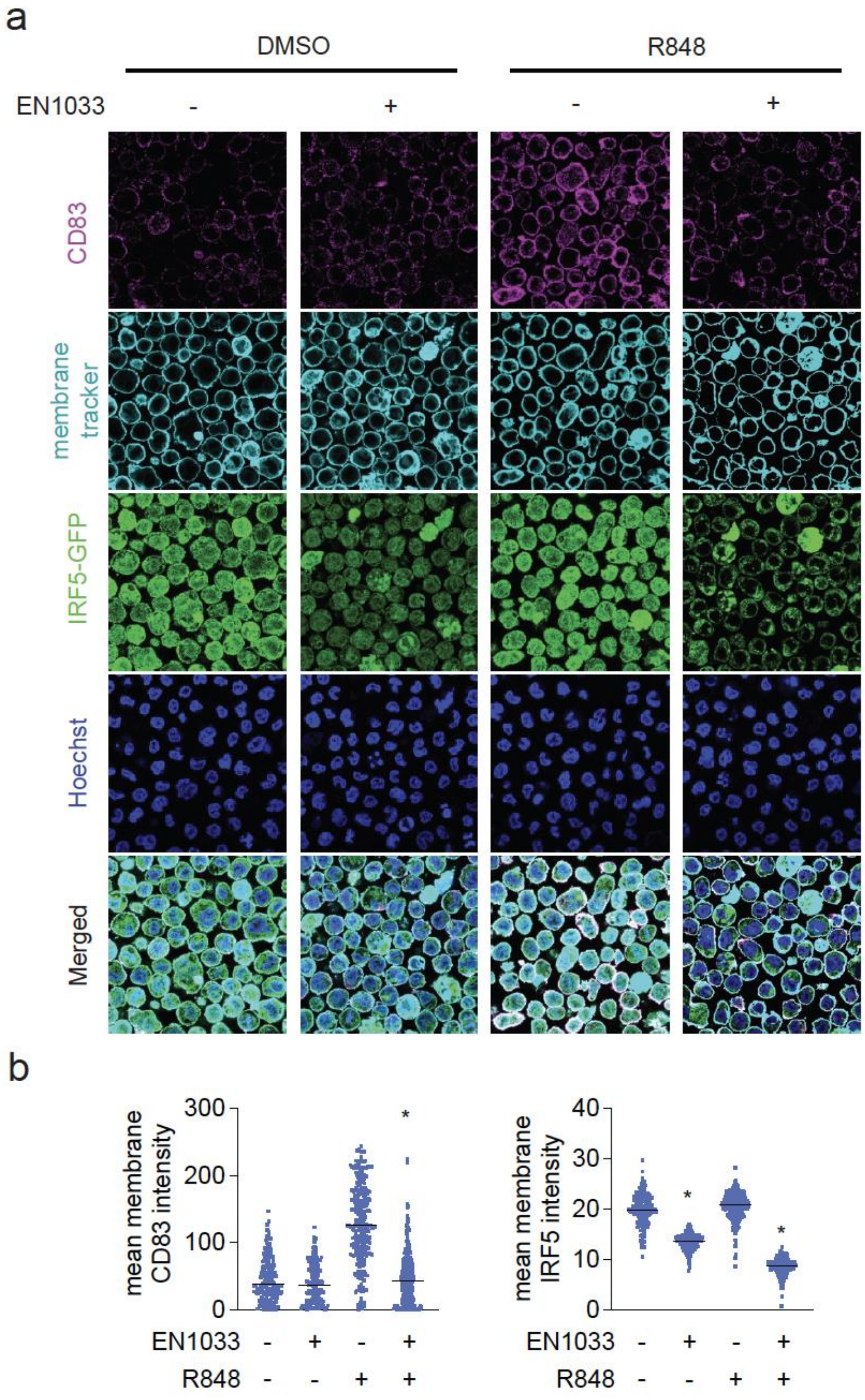


**Figure S7. Immunofluorescence staining of CD38 and IRF5 in THP1 cells. (a,b)** IFF5-GFP THP1 cells were activated with DMSO vehicle or R848 (10 μM) for 1 hour and then treated with DMSO vehicle or EN1033 (50 μM) for 24 hours. CD83, cell membrane and nucleus were then stained and imaged by confocal microscopy (a). Mean intensity of CD83 and IRF5 signal per cell is quantified and graphed in (b) shows individual values with average.

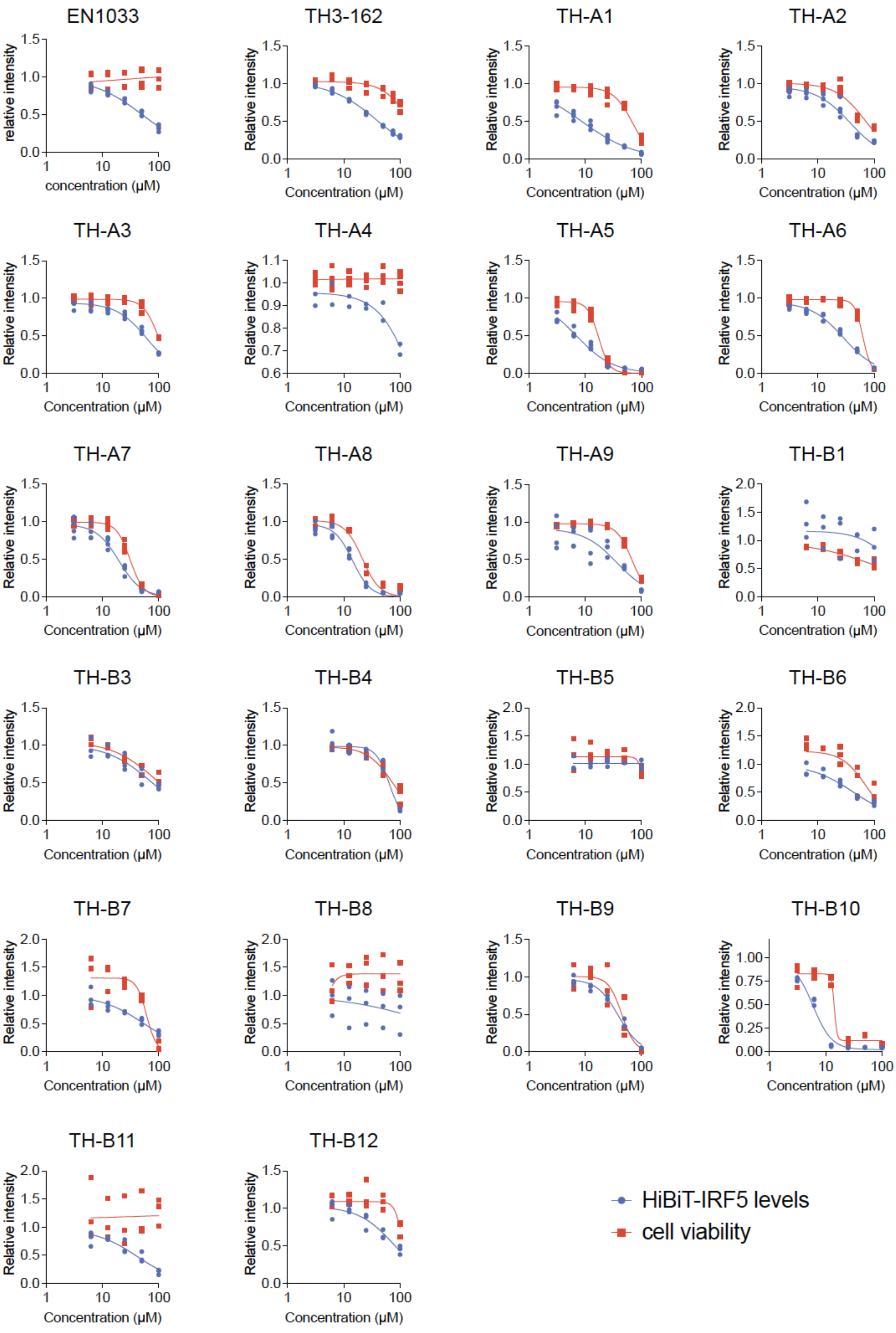


**Figure S8. Dose-response of HiBiT-IRF5 degradation with TH-B10 analogs.** HiBiT-IRF5 THP1 cells were treated with DMSO vehicle or compounds for 24 h, and HiBiT-IRF5 levels and cell viability were detected by luminescence. Data shown are individual replicate values from n=3 biologically independent replicates/group.

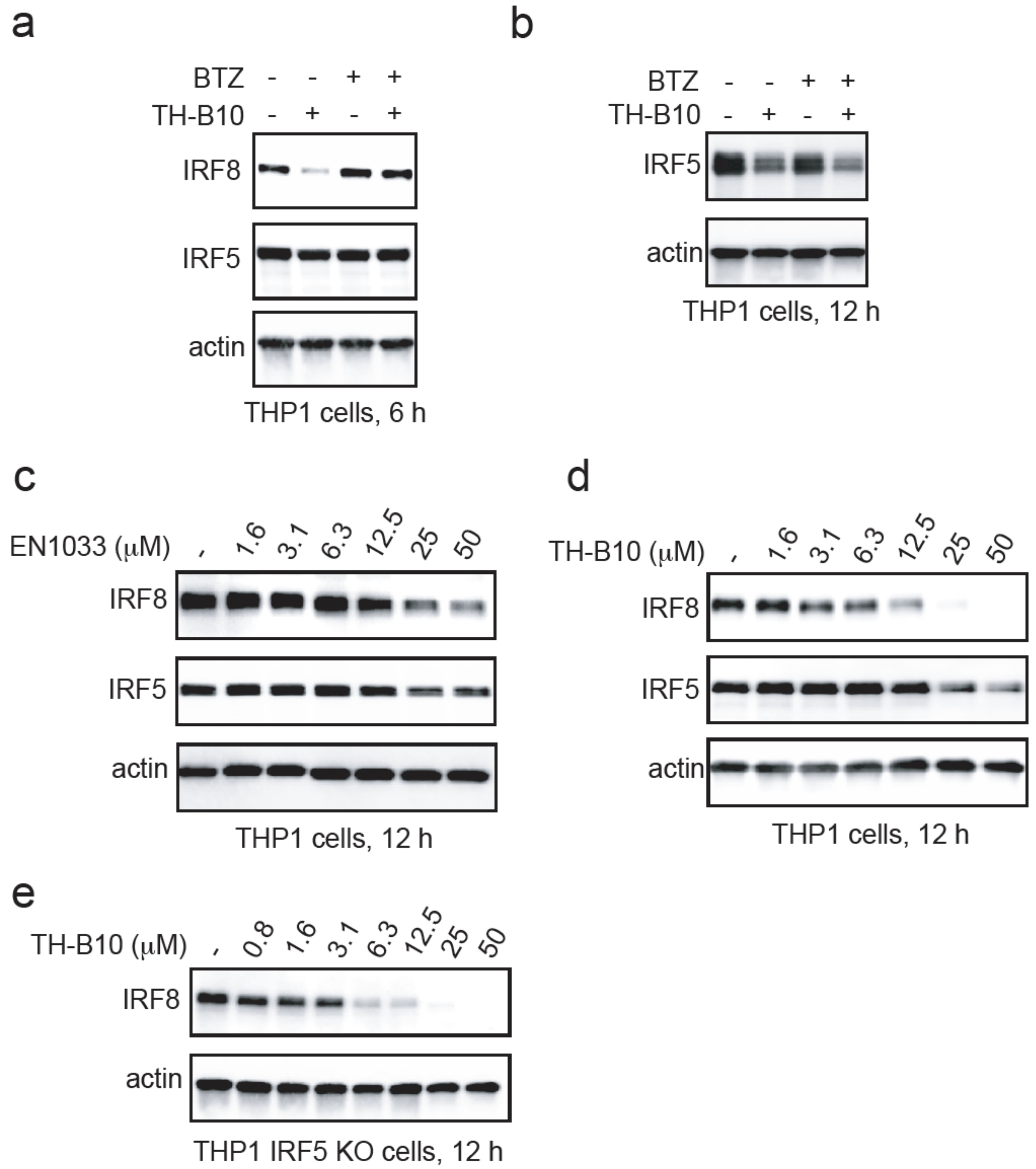


**Figure S9. Further characterization of IRF8 and IRF5 suppression by TH-B10. (a)** Proteasome-dependence of IRF8 degradation. THP1 cells were pre-treated with DMSO vehicle or BTZ (500 nM) 1 h prior to treatment with DMSO vehicle or TH-B10 (10 μM) for 6 h, after which IRF8, IRF5 and loading control actin levels were assessed by SDS/PAGE and Western blotting.(b) Proteasome-independence of IRF5 suppression. THP1 cells were pre-treated with DMSO vehicle or BTZ (500 nM) 1 h prior to treatment with DMSO vehicle or TH-B10 (10 μM) for 12 h, after which IRF5 and loading control actin levels were assessed by SDS/PAGE and Western blotting. **(c,d, e)** Dose-response of IRF8 and IRF5 degradation. THP1 cells were treated with DMSO vehicle or EN1033 **(c)** or TH-B10 **(d)** for 12 h, after which IRF8, IRF5, and loading control actin levels were assessed by SDS/PAGE and Western blotting. IRF5 KO THP1 cells were treated with DMSO vehicle or TH-B10 for 12 h, after which IRF8 and loading control actin levels were assessed by SDS/PAGE and Western blotting. Plots in figure **(a,b,c,d,e)** are representative of n=3 biological independent replicates.

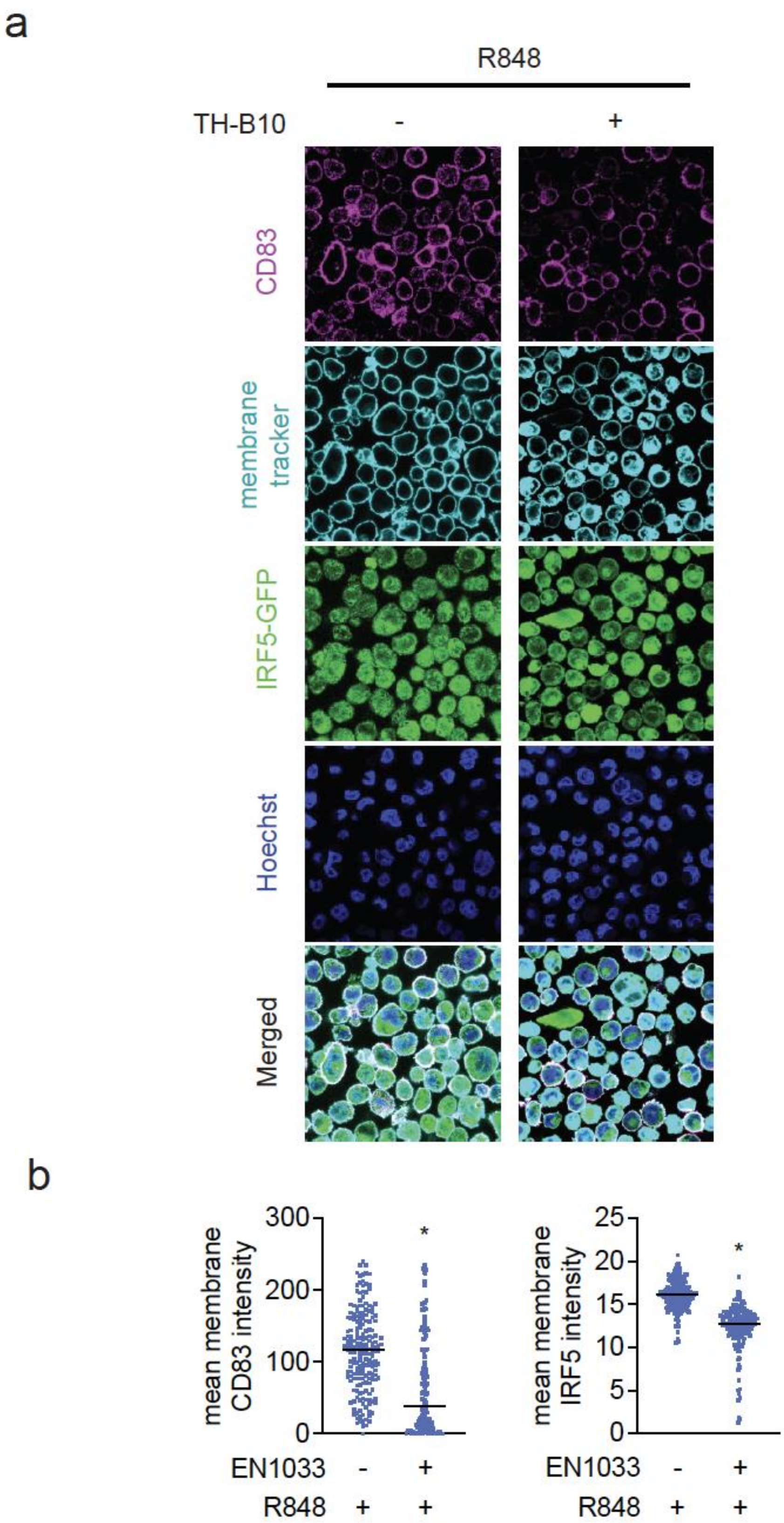


**Figure S10. Immunofluorescence staining of CD38 and IRF5 in activated THP1 cells after TH-B10 treatment. (a)** IFF5-GFP THP1 cells were activated with R848 (10 µM) for 1 hour and then treated with DMSO vehicle or TH-B10 (10 µM) for 12 hours. CD83, cell membrane and nucleus were then stained and imaged by confocal microscopy **(a)**. Mean intensity of CD83 and IRF5 signal per cell is quantified and graphed in **(b)** shows individual values with average.

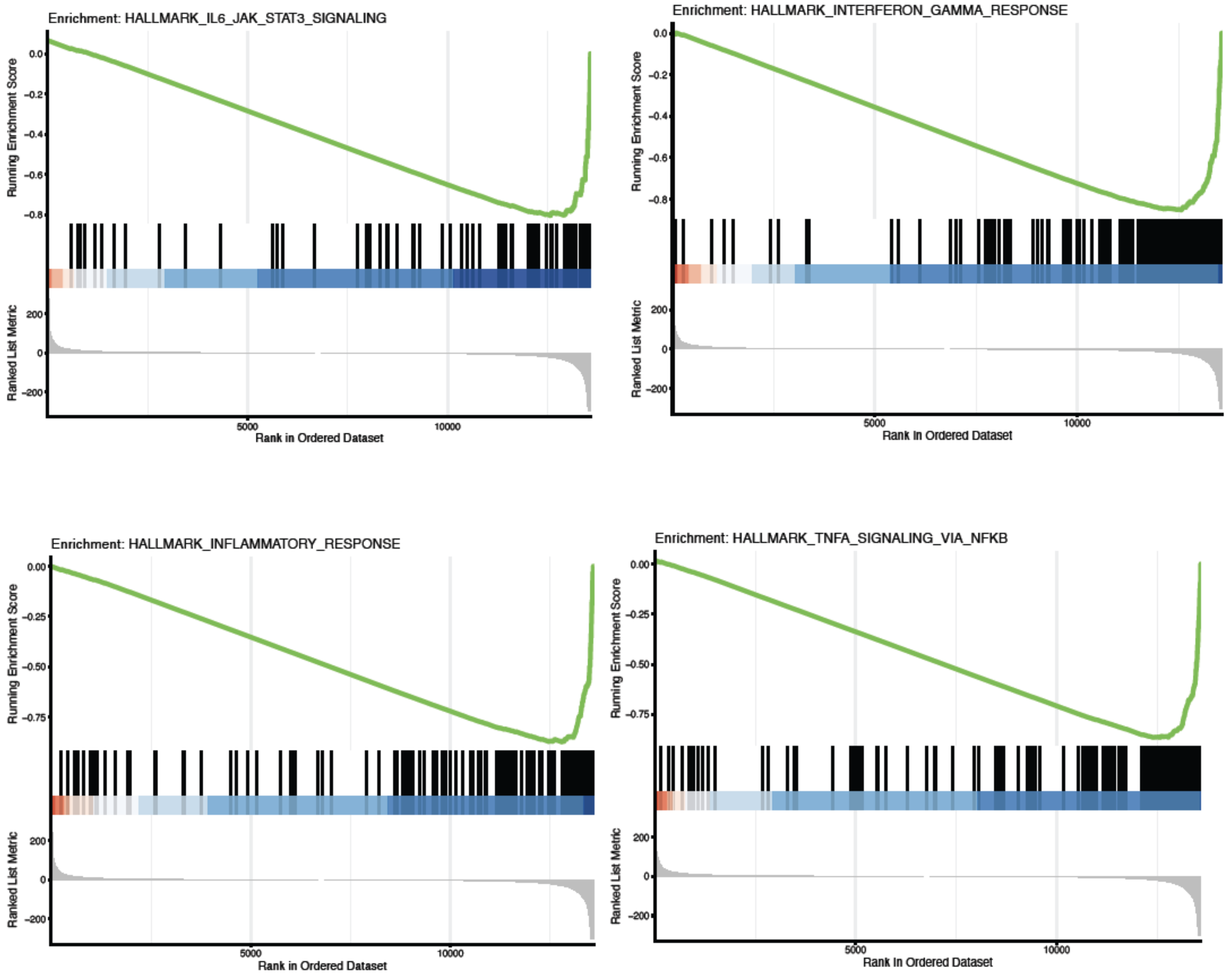


**Figure S11.** Enrichment plots for IRF5/IRF8 target genes from RNA seq analysis described in **Table S7** from THP1 cells treated with DMSO vehicle or TH-B10 (10 μM) for 12 h.

**Supporting Methods**

**Cell Culture**. THP1 cells were obtained from UC Berkeley's Biosciences Divisional Services Cell Culture Facility. THP1-dual cells, THP1-dual KO-IRF5 cells were purchased from Invivogen. THP1 based cells were cultured in RPMI media, and HEK293T cells were cultured in DMEM base media. Both media types contained 10% (v/v) fetal bovine serum (FBS), were supplemented with 1% glutamine, and were maintained at 37 °C with 5% $CO_2$.

**Covalent Ligand Library and Synthesis of Other Compounds**. Covalent ligands starting with "EN" are commercially available from Enamine LLC and compounds starting with "THA", "THB" are customized at Enamine LLC. Syntheses of other compounds are described in Supporting Information.

**Hibit IRF5 Cell Line Generation.** IRF5 construct was purchased from Genscript (cat. OHu27931). IRF5 sequence (without start codon) was then subcloned into a second-generation lentiviral transfer plasmid under control of the human UBC promoter fused to an N-terminal HiBiT (MGSVSGWRLFKKIS) tag [PMID: 28892606] and a 10 amino acid GS linker (GGGGSGGGGS). HEK293T cells were seeded into a 6-well plate in DMEM with 10% qualified FBS. The next day, cells were transfected with second-generation envelope (pMD2.G: Addgene #12259), packaging plasmids (psPAX2: Addgene #12260), and lentiviral transfer plasmid using Lipofectamine 3000 (Thermo Fisher #L3000008). Lentivirus-containing culture supernatant was harvested two days after transfection, passed through a 0.45 µm PES filter (CELLTREAT #229749), and added to cultures of THP1 cells at a final dilution of 1:9. Cells were selected with 1 µg/mL puromycin starting two days following transduction and for 14 days.

**Covalent Ligand Screen with IRF5 HiBiT Cell Line**. Covalent ligand screen and dose responses were conducted using a Promega Nano-Glo HiBiT Lytic Detection System (N3040) and CellTiter-Glo 2.0 Assay (G9242). HiBiT cells were seeded into 96-well plates (Corning 3917) at $1e^5$ cells per 100 µL of media. Cells were treated with 50 µL of media containing 50 µM compounds and treated for 24 h. The lytic detection system recipe was followed per Promega's suggestion. 50 µL of Lytic detection system reagents or CTG was added to each well. Plates were rocked for 15 min prior to their luminescence readout on the Tecan Spark Plate reader (30086376).

**Cell Viability Assay**. Cells were seeded in 96-well white plates then treated with DMSO vehicle control or compounds and incubated at 37 °C for 24 h. Cell viability assay was performed using CellTiter-Glo 2.0 reagent (Promega, G9241) according to manufacturer's protocol. Luminescent signals were measured using the Tecan Spark Plate reader (30086376).

**Western Blotting**. Pelleted cells are lysed with RIPA lysis buffer containing protease inhibitor cocktail (Pierce A32955). Samples were centrifuged at 20,000g for 20 min at 4 °C to remove cell debris. Samples' protein content was normalized to equal concentration. Samples were then boiled for 5 min at 95 °C after addition of 4× reducing Laemmli SDS sample loading buffer (Alfa Aesar) and ran on precast 4−20% Criterion TGX gels (Bio-Rad). Antibodies to IRF5 (Abcam, AB231163), IRF8 (Abcam, AB251475) beta-Actin (Sigma Aldrich, SAB1305546), GAPDH (Cell Signaling Technology, 14C10), were diluted as per recommended manufacturers' procedures. Proteins were resolved by SDS/PAGE and transferred to nitrocellulose membranes using the BioRad system (1704271 and 1704150). Blots were blocked with 5% BSA in Tris-buffered saline containing Tween 20 (TBST) solution for 1 h at room temperature, and probed with primary antibody diluted in recommended diluent per manufacturer overnight at 4 °C. Following washes with TBST, the blots were incubated in the dark with secondary antibodies purchased from Ly-Cor and used at 1: 5000 dilutions in 5% BSA in TBST at room temperature. Blots were visualized using an Odyssey LiCor scanner after additional washes. If additional primary antibody incubations were required, the membrane was stripped using ReBlot Plus Strong Antibody Stripping Solution (EMD Millipore, 2504), washed, and blocked again before being reincubated with primary antibody.

**Bortezomib, TAK243 or BafA1 Rescue Studies**. For 96-well plate format, HiBiT cells were plated at 100,000 cells per 100µL of media per well. Cells were pretreated for 1 h with media containing 1 µM Bafilomycin A1 (Medchemexpress, HY-100558), 0.5 µM Bortezomib (Cayman, C835F70) or 1 µM TAK243 (Ambeed, 1450833-55-2). Subsequently, EN1033 or DMSO control was added for a final concentration of 50 µM. Cells were then incubated at 37 °C, 5% $CO_2$ for 24 h. 100 µL lytic detection system reagent was then added to each well. Plates were rocked for 10 minutes prior to their luminescence read on the Tecan Spark Plate reader (30086376).

For Western blots, 1e$^{6}$ cells per 1mL of media were plated in 6-well plates. Cells were pretreated for 1 h with either 1 µM Bafilomycin A1 (Medchemexpress, HY-100558), 0.5 µM Bortezomib (Cayman, C835F70) or 1µM MLN4924 (Tocris Bioscience, 649910). Cells were then treated with EN1033 (50µM) or THB10 (10 µM) or DMSO until desired time point. Cells were harvested and assessed via western blot.

**IRF5 protein expression and purification.** IRF5 (without start codon) was sub-cloned into a standard bacterial cloning vector under control of an 8xTetOn dox-inducible promoter with an N-terminal TEV protease cleavable 6x-HisTag (MGSHHHHHHENLYFQS) and the human growth hormone (hGH) polyA terminator. 30 million HEK293T cells engineered to express reverse tetracycline transactivator (rTTA) were transfected in bulk with Lipofectamine 3000 and plated in 10 T225 flasks. 4 hours after plating cells were treated with 2uM doxycycline and incubated for 2-3 days until confluent. Cells were washed and collected with PBS and pelleted at 150 RCF for 3 minutes. Add 1mL per 0.05g of cell pellet of Mammalian Cell Lysis Buffer [GoldBio # GB-180] with 1:100 HALT Protease inhibitor Cocktail [Thermo Fisher # 78429] and incubate on ice for 15-30 minutes. Clarify at 20k RCF for 15 min at 4 °C and purify on a 1 mL nickel column. Wash and elution buffers were prepared by adding 10% glycerol (w/v) and 50 or 300 mM buffered imidazole to PBS. Elution was dialyzed in PBS + 10% glycerol (w/v).

**Covalent Ligand Screen and Gel-Based ABPP with IRF5/ IRF8 Pure Protein**. IRF5 pure protein or IRF8 (Origene, TP317646) (0.1 µg/25 µL in PBS) was treated with either DMSO vehicle or covalent ligand at 37 °C for 30 min and subsequently treated with 0.1 µM IA-rhodamine (Setareh Biotech) for 1 h at RT in the dark. The reaction was stopped by the addition of 4× reducing Laemmli SDS sample loading buffer (Alfa Aesar). After boiling at 95 °C for 5 min, the samples were separated on precast 4−20% Criterion TGX gels (Bio-Rad) and were analyzed by in-gel fluorescence using a ChemiDoc MP (Bio-Rad).

**Mapping of EN-1033 Site of Modification on IRF5 and IRF8 by LC-MS/MS**. Pure IRF5 or IRF8 protein was diluted in PBS (100 µL) and preincubated with EN1033 (100 µM final concentration) for 1 h at room temperature. The protein was precipitated by the addition of acetone (80% v/v) and incubation at −20 °C for 2 h. The sample was then spun at 20,000g for 10 min at 4 °C. The supernatant was carefully removed, and the sample was washed three times with 200 µL of ice-cold 0.01 M HCl/90% acetone solution, with spinning at 20,000 for 5 min at 4 °C between washes. The sample was then resuspended in 30 µL of 8 M urea in PBS and 30 µL of ProteaseMax surfactant (20 µg/mL in 100 mM ammonium bicarbonate, Promega, V2071) with vortexing. Ammonium bicarbonate (40 µL, 100 mM) was then added for a final volume of 100 µL. The sample was reduced with 10 µL of TCEP (10 mM final concentration) for 30 min at 60 °C and alkylated with 10 µL of iodoacetamide (12.5 mM final concentration) for 30 min at 37 °C. The sample was then diluted with 120 µL of PBS before 1.2 µL of ProteaseMax surfactant (0.1 mg mL−1 in 100 mM ammonium bicarbonate, Promega, V2071) and sequencing grade trypsin (10 µL, 0.5 mg mL−1 in 50 mM ammonium bicarbonate, Promega, V5111) were added for overnight incubation at 37°C. The next day, the sample was acidified with formic acid (5% final concentration) and fractionated using high pH reversed-phase peptide fractionation kits (Thermo Fisher, 84868) following manufacturer's protocol.

**TH3-189 Probe Labeling on IRF5 and IRF8 Pure Protein**. IRF5 or IRF8 pure protein (0.2 µg/50 µL in PBS) was treated with either DMSO vehicle or TH3-189 at 37 °C for 1 h. Click reagent Azide-Fluor 545 (Click Chemistry Tools, Inc. AZ109− 5), copper(II) sulfate, and TBTA (TCI Chemicals, T2993) were added to have a final concentration of 21.8 µM, 873.4 µM, and 47.2 µg/mL, respectively, for 1 h at RT in the dark. The reaction was stopped by addition of 4× reducing Laemmli SDS sample loading buffer (Alfa Aesar). After boiling samples at 95 °C for 5 min, the samples were separated on precast 4−20% Criterion TGX gels (Bio-Rad). Probe-labeled proteins were analyzed by in-gel fluorescence using a ChemiDoc MP instrument (BioRad).

**Luciferase Reporter Assay**. THP1 dual reporter or IRF5 KO dual reporter cells (Invivogen) were seeded at a density of 1e$^{6}$ cells/well in 96-well white plate. In a volume of 100 µL each well, the cells are treated with R848 (10 µM) or IFNa (10000 U/mL), and dose of compounds in 0.5 % FBS, RPMI glutamax media for 24 h. Lucia or Gaussia luciferase reporter genes' expression levels were examined by QUANTI-Luc$^{tm}$ 4 Lucia/Gaussia kit (Invivogen, rep-qlc4lg1) according to manufacturer's protocols. Luminescent signals were measured using the Tecan Spark Plate reader (30086376) with 1000 ms/well integration time.

**Cellular Thermal Shift Assay (CETSA).** THP1 cells were seeded at a density of $1e^7$ cells per 10 mL of media and were treated with either DMSO vehicle control or EN1033 (100μM) for 4 h or THB10 (50 μM) for 1h. Cells were harvested and washed twice with PBS, then suspended in 1 mL of PBS containing protease inhibitor cocktail (Pierce A32955). The cell suspension was then lysed, and the protein concentration was then normalized to equal concentration. The cell lysates were then aliquoted into eight 0.2 mL PCR tubes, each with a 100 μL volume. PCR strips were subjected to a temperature gradient via a program on Bio-Rad's T100 Thermal cycler. Samples were heated at their respective temperatures for 3 min, then at 25 °C for 3 min. Afterwards, cells were immediately snap-lysed in liquid nitrogen (3 freeze-thaw cycles). Cell debris along with precipitated and aggregated proteins were removed by centrifuging samples at 20,000 g for 20 min at 4 °C. 80μL of centrifuged samples were transferred to new PCR tubes and boiled for 5 min at 95 °C after addition of 4×reducing Laemmli SDS sample loading buffer (Alfa Aesar). Samples were analyzed by Western Blot analysis.

**IRF8 Knockdown Studies**. MISSION shRNA lentiviral construct, pMD2.G (Addgene, 12259) and psPAX2 (Addgene, 12260) were transfected into HEK293T cells using Lipofectamine 2000 (ThermoFisher, 11668027). The virus-containing medium was collected and filtered after 48 h and was used to infect THP1 target cells with a 1:1000 dilution of Polybrene (Sigma-Adrich, TR-1003-G). After 48 h, the infected cells were selected with puromycin. MISSION pLKO.1-puro non-mammalian shRNA Control (Sigma-Adrich, SHC016) was used as a control shRNA. The shRNA sequence used for generation of IRF8 knockdown lines is shown below. shIRF8 (Sigma-Aldrich, TRCN0000020984): GCCCGCATCATGATTAAAGAA.

**Generating Cells Expressing FLAG-Wild-Type or Mutant Human IRF5 or IRF8**. Wild type IRF5 sequences were subcloned with an N-terminal FLAG tag into a second-generation lentiviral transfer plasmid under control of an 8xTetOn promoter expressing in tandem reverse tetracycline transactivator (rTTA) under control of the strong constitutive EF1A promoter. Cysteine mutants of the original WT IRF5 sequence were generated using PCR. IRF8 WT and mutant constructs were customized and purchased from GenScript. These constructs were then used to generate stable FLAG-IRF5 or FLAG-IRF8-expressing cells using lentiviral infection. Before transfection, HEK293T cells were conditioned in complete media containing heat-inactivated FBS. For lentivirus production, FLAG-tagged wild type IRF5/IRF8 or FLAG-tagged IRF5/IRF8 mutant plasmids, psPAX2 (Addgene, 12260) and, pMD2.G (Addgene, 12259) were transfected into HEK293T cells using Lipofectamine 2000 (ThermoFisher, 11668027) in Opti-MEM (Gibco, 31985062). The virus-containing medium was collected and filtered (0.45 μm PES) after 48 hours. The virus was then used to infect new HEK293T with a 1:1000 dilution of Polybrene (Sigma-Aldrich, TR-1003-G). After 48 hours, infected cells were selected with puromycin (Abcam, ab141453) (2 μg/mL for HEK293T cells. After 3 days of the selection, cells were recovered by replacing puromycin-containing media with fresh media.

**Stress Granule Assessment**. HEK293T cells were seeded in 6-well plates at 500 000 cells per well (2 mL total) and left to adhere overnight. The cells were then transfected with the pEGFPG3BP1-WT plasmid (Addgene, #135997). For plasmid transfection, 0.75 μg of overexpression plasmid was added to each well together with the Lipofectamine 3000 transfection reagent and P3000 (Invitrogen, L3000001) in accordance with the manufacturer's protocol. After 48 h, the cells were reseeded in an 8-well glass bottom cell culture dish (Ibidi, 80807). After 12 h, the adherent cells were then treated with DMSO (vehicle), $NaAsO_2$ or EN1033 for 8 h. The cells were then fixed with 4% paraformaldehyde, methanol-free (Cell Signaling, #47746) for 15 min, washed with PBS three times, and then stained with 1 μM Hoechst stain (Invitrogen, H3570), with all steps performed at room temperature. Images were captured using a Zeiss LSM880 FCS microscope with a 40× oil objective.

**Quantification of Cytokines**. THP1 cells were treated with R848 (10 μM) and EN1033 (100 μM) or TH3-116 (100 μM) for 20 h. Quantification of TNF$\alpha$, IL-6 and CD83 were determined by ELISA using individual kits following the manufacturer's instructions (e.g. Human IL-6 ELISA kit (EH2IL6) from Invitrogen, Human TNF$\alpha$ ultrasensitive ELISA kit (KHC3014) from Invitrogen, Human CD83 simplestep ELISA kit (ab277711) from Abcam). ELISAs were read using a Tecan Spark Plate reader (30086376).

**Immunofluorescence.** To detect CD83 presentation on THP1 cell surface, THP1 cells (IRF5-GFP) were pretreated with R848 agonist (10 μM) or DMSO for 1h. After that, EN1033 (50 μM) or THB10 (10 μM) was introduced for 24 h or 12 h, respectively. The cells were then fixed with 4% paraformaldehyde, methanol-free (Cell Signaling, #47746) for 15 min, permeabilized with 0.1% Triton in PBS for 15 min at room temperature. Cells were then washed with PBS and then blocked with 5% BSA in TBST for 1 h. CD83 primary antibody was diluted at suggested ratio with blocking buffer and incubated with cells over night at 4 ˚C. After washing with PBS three times, cells were incubated with Alexa Fluor-labeled 647 secondary antibodies for an additional 1 h. Cells were then washed with PBS three times and stained with 1 µM Hoechst (Invitrogen, H3570) for nucleus staining, cell mask (Invitrogen, C10045) for cell membrane staining. Confocal images were captured using a Zeiss LSM880 FCS microscope with a 40× oil objective. Image processing and analysis were performed using a Python script with nucleus and cell mask channels as parameters for detection of single cells, cell membrane and cytosol area.

**Covalent Ligand/Probes pulldown**. For Western Blotting Detection, THP1 cells were seeded at $2e^6$ cells/mL in flask (5 mL per replicate). Cells were then treated with compounds at 100 µM final concentration and incubated for 4 h. Pelleted cells were lysed by probe sonication on ice in PBS containing a protease inhibitor cocktail (Pierce A32955). Lysate protein concentrations were measured using BCA Protein Assay (Pierce 23225) and normalized to 5 mg/mL in 500 µL of PBS with protease inhibitor cocktail. A master mix of the click reagents was prepared such that each replicate would receive 10 µL of biotin picolyl azide (10 mM stock in DMSO, Sigma-Aldrich, 900912), 10 µL of copper (II) sulfate (50 mM stock in water, Sigma-Aldrich, 203165), 30 µL of TBTA (1.7 mM in 4:1 tBuOH/DMSO, TCI Chemicals, T2993) and 10 µL of TCEP (50mM in water freshly prepared, ThermoScientific, 20491). Samples were vortexed and incubated on a rotator at room temperature for 1 hour. 500 µL of cold methanol was added, and samples underwent three cold methanol washes with centrifugation to pellet proteins and sonication for resuspension. Pelleted proteins were redissolved in 200 µL of PBS with 1.2% SDS (w/v) by sonication. Samples were heated to 90°C for 5 minutes, and 5µL of the sample was saved for input (diluted to 60 µL and 20 µL of 4x reducing Laemmli SDS sample loading buffer). 55 µL (per sample) of streptavidin-agarose beads (ThermoFisher, 20353) were washed with PBS using a Micro-Bio Spin Column (Bio-Rad, 7326204) and vacuum manifold. 500 µL of PBS was added to the dissolved proteins, and then the washed beads were added to each sample using two washes of 250 µL of PBS. Samples were incubated on a rotator at 4°C overnight. Samples were then put in a 37°C bath to redissolve SDS for 5 minutes, and then beads were pelleted by centrifugation at 1400 g for 5 min. The supernatant was removed, and beads were washed with 0.2% SDS in PBS (w/v) for 10 minutes on a rotator at room temperature. The supernatant was removed, and pelleted beads were moved to Micro-Bio Spin Columns using two washes of PBS. Beads were washed on a vacuum manifold three times with PBS and then three times with water. The beads were then transfered to screw cap Eppendorf tubes with PBS, centrifuged, and the supernatant removed. 30 µL of 1x reducing Laemmli SDS sample loading buffer was added to each sample and then boiled at 95 °C for 15 minutes to elute proteins from beads. Samples and corresponding inputs were run following Western Blot procedure.

**Pulldown TMT Proteomics Experiment with Covalent Ligand/Probes**. THP1 cells were seeded at $2e^6$ cells/mL in flask (10 mL per replicate). Cells were then treated with EN1033 compound at 100 µM final concentration and incubated for an additional 4 h. Cells were then collected and washed twice with PBS. Cells were lysed by probe sonication on ice in PBS containing a protease inhibitor cocktail (Pierce A32955). Lysate protein concentrations were measured using BCA Protein Assay (Pierce 23225) and normalized. A master mix of the click reagents was prepared such that each replicate would have 10 µL of biotin picolyl azide (10 mM stock in DMSO, Sigma-Aldrich, 900912), 10 µL of copper (II) sulfate (50 mM stock in water, Sigma-Aldrich, 203165), 30 µL of TBTA (1.7 mM in 4:1 tBuOH/DMSO, TCI Chemicals, T2993) and 10 µL of TCEP (50 mM in water freshly prepared, ThermoScientific, 20491). Samples were vortexed and incubated on a rotator at room temperature for 1 hour. To each sample, 100% acetonitrile was added to the final concentration of 80% and mixed by inversion to precipitate proteins. Proteins were pelleted by centrifugation at 21,000 g at 4 °C for 10 minutes. The supernatant was removed, 500 µL of cold methanol was added, and samples underwent three cold methanol washes with centrifugation to pellet proteins and sonication for resuspension. After the final wash, the pellets were left to air dry for 10 minutes. Proteins were redissolved in 1 mL of PBS with 1.2% SDS (w/v) by sonication. Samples were then heated to 90°C for 5 minutes, and 5 mL of PBS was transferred to 15 mL tubes. 170 µL of streptavidin-agarose beads were added to each sample and incubated at 4°C overnight with constant rotation. Samples were then put in a 37°C bath to redissolve SDS for 5 minutes, and then beads were pelleted by centrifugation at 1400 g for 5 min. The supernatant was removed, and beads were washed with 0.2% SDS in PBS (w/v) for 10 minutes on a rotator at room temperature. The supernatant was removed, and pelleted beads were transferred to Micro-Bio Spin Columns using two washes of PBS. Beads were washed on a vacuum manifold four times with PBS and then four times with DI water. The beads were then transferred to screw cap Eppendorf tubes with two 250 µL washes of 6 M Urea in PBS. To each sample, 25 µL of DTT (30 mg/mL in water) was added and incubated at 65 °C for 20 minutes, gently mixing every 5 minutes. Samples were cooled to room temperature, and then 25 µL of iodoacetamide (400 mM in water) was added and incubated at 37 °C for 30 minutes with constant agitation. Samples were then diluted with PBS, centrifuged, the supernatant removed, rewashed with PBS, and then washed once with 500 µL of 50 mM TEAB (ThermoScientific 90114), and the supernatant removed. Beads were resuspended in 100 µL of 50 mM TEAB, and 4 µL of sequencing grade trypsin (0.5 mg/mL, Promega, V5111) was added to each sample. Samples were digested at 37°C overnight with constant agitation. Samples (beads and liquid) were then moved to a Micro-Bio Spin Column and centrifuged at 2000 g for 2 min at RT to collect flow through. Quantification of peptides was done using the Thermo Scientific Pierce Quantitative Colorimetric Peptide Assay (ThermoScientific, 23275). 100 µL of each sample was for TMT labeling. TMT labels (ThermoScientific, 90061, or A58332) were equilibrated with anhydrous acetonitrile, and 20 µL of each tag was added to a corresponding replicate. The reaction was incubated at room temperature for 1 hour on a rotating mixer and then quenched with 5% hydroxylamine for 15 minutes at room temperature. Peptides were dried down by Vacufuge (Eppendorf, 022820168) at 30 °C until dry. Samples were then resuspended in 50 µL of 0.1% Trifluoroacetic acid (TFA) in water and combined. Using the previously recorded concentrations, 100 µg of peptides were then fractionated according to the manufacturer's protocol using the Thermo Fisher High pH Reverse Phase Fractionation Kit (ThermoScientific, 84868). Following fractionation, samples were dried down using Vacufuge at 30°C and then resuspended in 25 µL of 0.1% formic acid in water by vortexing and bath sonication. Samples were centrifuged at 20,000 g for 10 minutes at 4 °C and the supernatant were transferred into an LCMS vial with a glass insert (ThermoScientific, 6PME03C1SP) and capped. Samples were analyzed by LC-MS/MS.

**Isotopic Desthiobiotin (isoDTB)-ABPP Cysteine Chemoproteomic Profiling of EN-1033 and THB10**. THP1 cells were treated with either EN-1033 (100 µM) or DMSO for 6 h before cell collection and lysis by probe sonication. For THB10, THP1 cells were treated with THB10 (10 µM) or DMSO for 1h before collection and lysis by pro sonication. The proteome concentrations were determined using BCA assay and adjusted to 2 mg/mL. For each biological replicate, 2 aliquots of 1 mL of 2 mg/mL were used (i.e., 4 mg per condition). Each aliquot was treated with 20 µL of IA-alkyne (26.6 mg/mL in DMSO, 200 µM final concentration) for 1 h at RT. Two master mixes of the click reagents were prepared in the meanwhile, each containing 510 µL TBTA (0.9 mg/mL in 4:1 tBuOH/DMSO), 165 µL $CuSO_4$ (12.5 mg/mL in H2O), 165 µL TCEP (14.0 mg/mL in H2O) and 160 µL of either heavy or light isoDTB tags (4 mg in DMSO, Click Chemistry Tools, 1565). The samples were then treated with 120 µL of the heavy (DMSO treated) or light (compound treated) master mix for 1 h at RT. After incubation, one light and one heavy labeled samples were combined and acetone-precipitated overnight at −20 °C. The samples were then centrifuged at 3,500 rpm for 10 min, acetone was removed, and the protein pellets resuspended in cold MeOH by sonication. The samples were centrifuged at 3,500 rpm for 10 min and MeOH was removed (repeated 3× in total). The pellets were dissolved in 600 µL urea (8 M in 0.1 M TEAB) by sonication and the urea concentration was then adjusted to 2 M by adding 1800 µL of TEAB (0.1 M). Two tubes containing solubilized proteins were combined, further diluted with 2400 µL 0.2% NP40 in PBS, and bound to highcapacity streptavidin agarose beads (200 µL/sample, ThermoFisher, 20357) for 1 h at RT with mixing. The beads were then centrifuged for 1 min at 1,000 g, the supernatant was removed, and the beads were washed 3 times with 0.1% NP40 in PBS, 3 times with PBS and 3 times with H2O. The samples were then resuspended in 8 M urea (600 µL in 0.1 M TEAB) and treated with DTT (30 µL, 31 mg/mL in H2O) for 45 min at 37 °C. They were then reacted with iodoacetamide (30 µL, 74 mg/ mL in H2O) for 30 min at RT, followed by DTT (30 µL, 31 mg/mL in H2O) for 30 min at RT. The samples were diluted with 1800 µL TEAB (0.1 M), centrifuged for 1 min at 1,000 g, and the supernatant was removed. The beads were resuspended in 400 µL urea (2 M in 0.1 M TEAB), and trypsin (8 µL, 0.5 mg/mL) was added and incubated for 20 h at 37 °C. The samples were then diluted with 800 µL 0.1% NP40 in PBS and the beads were washed 3 times with 0.1% NP40 in PBS, 3 times with PBS, and 3 times with H2O. Peptides were then eluted with 0.1% formic acid in 50% acetonitrile (3 × 400 µL). The samples were then dried using a vacuum concentrator at 30 °C, resuspended in 300 µL 0.1% TFA in H2O, and fractionated using high pH reversed-phase peptide fractionation kits (ThermoFisher, 84868) according to the manufacturer's protocol.

**IsoDTB-ABPP Mass Spectrometry Analysis**. Mass spectrometry analysis was performed on an Orbitrap Eclipse Tribrid Mass Spectrometer with a High Field Asymmetric Waveform Ion Mobility (FAIMS Pro) Interface (Thermo Scientific) with an UltiMate 3000 Nano Flow Rapid Separation LCnano System (Thermo Scientific). Off-line fractionated samples (5 µL aliquot of 15 µL sample) were injected via an autosampler (Thermo Scientific) onto a 5 µL sample loop which was subsequently eluted onto an Acclaim PepMap 100 C18 HPLC column (75 µm x 50 cm, nanoViper). Peptides were separated at a flow rate of 0.3 µL/min using the following gradient: 2% buffer B (100% acetonitrile with 0.1% formic acid) in buffer A (95:5 water:acetonitrile, 0.1% formic acid) for 5 min, followed by a gradient from 2 to 40% buffer B from 5 to 159 min, 40 to 95% buffer B from 159 to 160 min, holding at 95% B from 160 to 179 min, 95% to 2% buffer B from 179 to 180 min, and then 2% buffer B from 180 to 200 min. Voltage applied to the nano-LC electrospray ionization source was 2.1 kV. Data was acquired through an MS1 master scan (Orbitrap analysis, resolution 120,000, 400−1800 m/z, RF lens 30%, heated capillary temperature 250 °C) with dynamic exclusion enabled (repeat count 1, duration 60 s). Data-dependent data acquisition comprised a full MS1 scan followed by sequential MS2 scans based on 2 s cycle times. FAIMS compensation voltages (CV) of −35, −45, and −55 were applied. MS2 analysis consisted of: quadrupole isolation window of 0.7 m/z of precursor ion followed by higher energy collision dissociation (HCD) energy of 38% with an orbitrap resolution of 50,000. Data was extracted in the form of MS1 and MS2 files using Raw Converter (Scripps Research Institute) and searched against the Uniprot human database using ProLuCID search methodology in IP2 v.3-v.5 (Integrated Proteomics Applications, Inc.).37 Cysteine residues were searched with a static modification for carboxyaminomethylation (+57.02146) and up to two differential modifications for methionine oxidation and either the light or heavy isoDTB tags (+561.33872 or +567.34621, respectively). Peptides were required to be fully tryptic peptides. ProLuCID data were filtered through DTASelect to achieve a peptide false-positive rate below 5%. Only those probe-modified peptides that were evident across two out of three biological replicates were interpreted for their isotopic light to heavy ratios. Light versus heavy isotopic probe-modified peptide ratios are calculated by taking the mean of the ratios of each replicate paired light versus heavy precursor abundance for all peptide-spectral matches associated with a peptide. The paired abundances were also used to calculate a paired sample t test P value in an effort to estimate constancy in paired abundances and significance in change between treatment and control. P values were corrected using the Benjamini−Hochberg method.

**Quantitative TMT Proteomics Analysis**. THP1 cells were treated with either DMSO vehicle or EN-1033 or THB10 or DMSO for indicated time points and lysate was prepared by probe sonication. Briefly, 25−100 μg protein from each sample was then reduced, alkylated and tryptically digested overnight. Individual samples were then labeled with isobaric tags using commercially available TMTsixplex (Thermo Fisher Scientific, P/N 90061) kits, in accordance with the manufacturer's protocols. Tagged samples (20 μg per sample) were combined, dried using a vacuum concentrator at 30 °C, resuspended with 300 μL 0.1% TFA in H2O, and fractionated using high pH reversed-phase peptide fractionation kits (Thermo Fisher Scientific, P/N 84868) according to the manufacturer's protocol. Fractions were dried using a vacuum concentrator at 30 °C, resuspended with 50 μL 0.1% FA in H2O, and analyzed by LC-MS/MS as described below. Quantitative TMT-based proteomic analysis was performed as previously described using a Thermo Eclipse with FAIMS LC-MS/MS.5 Acquired MS data was processed using ProLuCID search methodology in IP2 v.3-v.5 (Integrated Proteomics Applications, Inc.). Trypsin cleavage specificity (cleavage at K, R except if followed by P) allowed for up to 2 missed cleavages. Carbamidomethylation of cysteine was set as a fixed modification, methionine oxidation, and TMTmodification of N-termini and lysine residues were set as variable modifications. Reporter ion ratio calculations were performed using summed abundances with the most confident centroid selected from the 20 ppm window. Only peptide-tospectrum matches that are unique assignments to a given identified protein within the total data set are considered for protein quantitation. High confidence protein identifications were reported with a < 1% false discovery rate (FDR) cutoff. Differential abundance significance was estimated using ANOVA with Benjamini-Hochberg correction to determine p-values.

**RNA Extraction and RNA Sequencing.** THP1 cells were seeded at $1e^6$ cells/mL, 2 mL each condition in flasks. The cells are then co-treated with R848 (10 μM) and DMSO or EN1033 (50 μM) for 24 hours or TH-B10 (10 μM) for 12 hours. After treatment, cells were washed twice with PBS before centrifuging to collect pellet cells. PBS supernatant was aspirated, and the cell pellet was further processed. The Monarch Total RNA Miniprep kit (New England Biolabs, T2010S) was used to harvest total RNA and was followed according to the manufacturer's protocols. RNA Sequencing Library preparation and sequencing was performed by the QB3-Berkeley Genomics core labs (cite). Total RNA quality, as well as poly-dT enriched mRNA quality, were assessed on an Agilent 2100 Bioanalyzer. Libraries were prepared using the KAPA mRNA Hyper Prep kit (Roche KK8581). Truncated universal stub adapters were ligated to cDNA fragments, which were then extended using 10 cycles of PCR using unique dual indexing primers into full length Illumina libraries. Library quality was checked on an AATI (now Agilent) Fragment Analyzer and transferred to the Vincent J. Coates Genomics Sequencing Laboratory (GSL), another QB3-Berkeley Core Research Facility at UC Berkeley.

**Analysis of RNAseq Data.** Differential expression analysis Samples were sequenced on one lane of a NovaSeq X Plus generating paired-end 150bp reads to a median depth of 59.3 million reads per sample (minimum 46.3

million reads). After demultiplexing, raw FASTQ files were pseudoaligned against the human reference transcriptome (Gencode v48) using kallisto with default parameters [1]. The median read assignment rate across all samples was 84.5% (minimum 81.5%). To test for differential expression, abundance estimates were summarized to the gene level using tximport and modeled using DESeq2 as a simple two-group comparison (treated vs control) [2,3]. Per-gene summary statistics including p-values were Benjamini-Hochberg adjusted and exported for downstream analysis.


**Synthetic Methods and Characterization** All chemical reactions were carried out under a nitrogen atmosphere with dry solvents under anhydrous conditions, unless otherwise noted. Reagents were purchased at the highest commercial quality and used without further purification, unless otherwise stated. Room temperature is defined as between 21-25 °C. Reactions were stirred magnetically and monitored by thin layer chromatography (TLC) using TLC plates precoated with silica gel 60 F254 on aluminium (Merck KGaA). Detection was by UV (254 nm and 365 nm) or chemical stain (KMnO4, ninhydrin, iodine). Solvents were removed in vacuo using either a Buchi R-300 Rotavapor (equipped with an I-300 Pro Interface, B-300 Base Heating Bath, Welch 2037B-01 DryFast pump, and VWR AD15R-40-V11B Circulating Bath). Solvents for silica gel chromatography were used as supplied by Sigma-Aldrich. Automated flash chromatography was performed on a Biotage Isolera instrument, equipped with a UV detector. Chromatograms were recorded at 254 and 280 nm. High-resolution mass spectra (HRMS) were obtained using Q ExactiveTM Plus Hybrid QuadrupoleOrbitrapTM Mass Spectrometer. $^{1}$H and $^{13}$C Nuclear Magnetic Resonance (NMR) spectra were recorded on BRUKER AV (600 MHz and 700 MHz), AVB (400 MHz), AVQ (400 MHz) and NEO (500 MHz) spectrometers. Measurements were carried out at ambient temperature. Chemical shifts (δ) are reported in ppm with the residual solvent signal as internal standard (chloroform at 7.26 and 77.2 ppm for $^{1}$H NMR and $^{13}$C NMR, respectively, methanol at 3.31 and 49.0, respectively and DMSO at 2.50 and 39.5, respectively). Multiplicity is reported as follows: singlet (s), doublet (d), doublet of doublet (dd) doublet of triplet (dt), triplet (t), triplet of doublet (td), quartet (q), and multiplet (m). Coupling constants (J) are reported in Hertz (Hz). 13C NMR spectra were recorded with broadband 1 H decoupling.

**Synthesis of 1-(2,3,11,11a-tetrahydro-1*H*-benzo[*e*]pyrrolo[1,2-*a*][1,4]diazepin-10(5*H*)-yl)propan-1-one (TH3-116)**

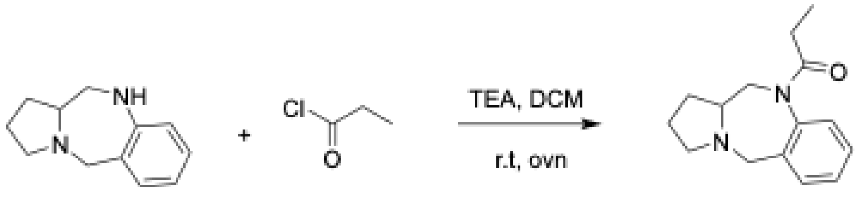


To a solution of 2,3,5,10,11,11a-hexahydro-1*H*-benzo[*e*]pyrrolo[1,2-*a*][1,4]diazepine (50 mg, 0.27 mmol) in DCM (5 mL) was added triethylamine (55 µL, 0.40 mmol) dropwise. The solution was stirred at 0 °C for 5 min. Propionoyl chloride (35 µL, 0.40 mmol) was added and the solution was warmed to room temperature. The reaction was then run over night. After completion of reaction. The product was extract with DCM and wash with brine and dry by $Na_2SO_4$. The crude residue was purified by silica gel chromatography (0-7% MeOH in DCM) to afford 42 mg (65%) of the title compound as a yellow-white oil. $^{1}$H NMR (500 MHz, $CDCl_3$) δ 7.31 (dd, *J* = 7.0, 2.2 Hz, 1H), 7.30 – 7.27 (m, 1H), 7.25 (d, *J* = 5.8 Hz, 1H), 7.16 (dd, *J* = 7.5, 1.6 Hz, 1H), 4.81 (dd, *J* = 13.3, 2.2 Hz, 1H), 3.76 (d, *J* = 13.4 Hz, 1H), 3.56 (d, *J* = 13.4 Hz, 1H), 3.10 (td, *J* = 8.6, 2.9 Hz, 1H), 2.64 (tdd, *J* = 9.3, 6.5, 2.1 Hz, 1H), 2.59 – 2.46 (m, 2H), 2.28 (dq, *J* = 15.2, 7.5 Hz, 1H), 2.03 – 1.94 (m, 2H), 1.86 – 1.71 (m, 2H), 1.45 (dddd, *J* = 12.5, 11.1, 9.6, 6.3 Hz, 1H), 1.05 (t, *J* = 7.5 Hz, 3H). $^{13}$C NMR (126 MHz, $CDCl_3$) δ 173.46,

143.17, 137.17, 131.03, 128.94, 128.47, 128.35, 77.71, 77.45, 77.20, 67.03, 57.74, 55.46, 53.86, 50.55, 28.87, 28.01, 21.53, 10.09. HRMS (ESI) m/z calcd for $C_{15}H_{20}N_2O^+$ [M+H]$^+$ : 245.1648; found: 245.1642

**Synthesis of 1-(7-ethynyl-2,3,11,11a-tetrahydro-1*H*-benzo[*e*]pyrrolo[1,2-*a*][1,4]diazepin-10(5*H*)-yl)prop-2-en-1-one (TH3-189)**

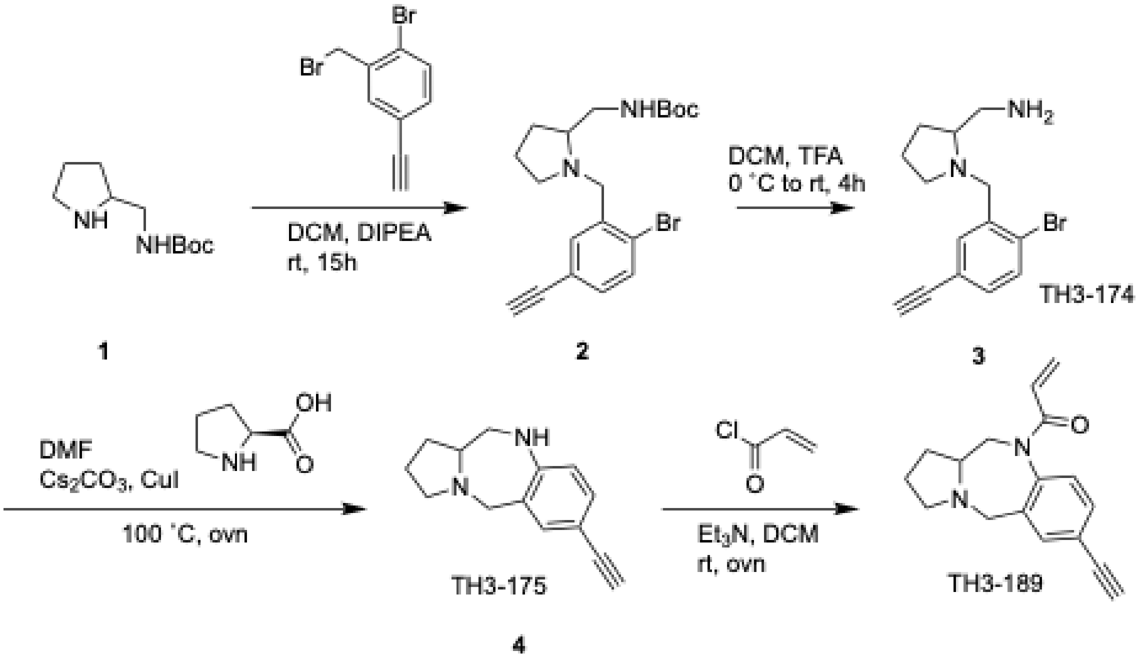


Step **1**: Tert-butyl (pyrrolidin-2-ylmethyl) carbamate (200mg, 1.0 mmol) was dissolved in DCM (5 mL) and DIPEA (2.5 mmol) was added at room temperature. After 30 min, solution of 1-bromo-2-(bromomethyl)-4-ethynylbenzene in DCM (2 mL) was added drop wise and continued stirring for 15 h. TLC and LCMS indicated the formation of alkylated product. After completion of the reaction, reaction mixture was diluted with ethyl acetate (30 mL) and washed with water. Next, the organic layer was separated, dried over $Na_2SO_4$ and purified using flash chromatography to obtain TH3-174 as colorless viscous liquid (88%). HRMS (ESI) m/z calcd for $C_{19}H_{26}BrN_2O_2^+$ [M+H]$^+$ : 393.1169; found: 393.1164.
Step **2**: TH3-174 (0.4 mmol) was dissolved in DCM (3 mL) and added TFA (0.5 mL) dropwise at 0°C and slowly warmed reaction mixture to room temperature and continued stirring for 4 h. After completion of the reaction, reaction mixture was concentrated to dryness, wash and extract the organic phase with DCM. The crude residue was purified using flash chromatography (0-10% MeOH in DCM). HRMS (ESI) m/z calcd for $C_{14}H_{18}BrN_2^+$ [M+H]$^+$ : 293.0645; found: 293.0618.
Step **3**: The obtained product from step 2 (80 mg, 0.11 mmol) was dissolved in DMF (2 mL). Next, $Cs_2CO_3$ (0.22 mmol), CuI (10 mol%) and L-proline (20 mol%) were added and the mixture was run at 100 °C for 15h. After completion of the reaction, reaction mixture was diluted with ethyl acetate (30 mL) and washed with cold water. Next, the organic layer was separated, dried over $Na_2SO_4$ and purified using flash chromatography (10% MeOH in DCM) to obtain compound TH3-175 as pale-yellow viscous liquid. $^1H$ NMR (500 MHz, $CDCl_3$) δ 7.26 (s, 1H), 7.20 (dd, *J* = 8.1, 1.9 Hz, 1H), 6.63 (d, *J* = 8.1 Hz, 1H), 3.79 (d, *J* = 13.7 Hz, 1H), 3.49 (d, *J* = 13.7 Hz, 1H), 3.35 (dd, *J* = 12.9, 2.3 Hz, 1H), 3.16 – 3.11 (m, 1H), 2.96 (s, 1H), 2.78 (dd, *J* = 12.9, 9.2 Hz, 1H), 2.53 – 2.47 (m, 2H), 1.95 – 1.76 (m, 3H), 1.49 (dddd, *J* = 12.1, 10.8, 9.6, 6.1 Hz, 1H).$^{13}C$ NMR (126 MHz, $CDCl_3$) δ 150.51, 134.80, 131.86, 128.97, 119.09, 113.70, 84.26, 77.45, 77.20, 76.95, 75.59, 68.07, 58.85, 55.94, 52.56, 29.08, 22.11.HRMS (ESI) m/z calcd for $C_{14}H_{17}N_2^+$ [M+H]$^+$ : 213.1385; found: 213.1368.
Step **4**: To a solution of TH3-175 (14 mg, 0.07 mmol) in DCM (5 mL) was added triethylamine (14 µL, 0.1 mmol) dropwise. The solution was stirred at 0°C for 20 min. prop2-enoyl chloride (8.5 uL, 0.1 mmol) was added and the solution was warmed to room temperature. The reaction was then run over night. After completion of reaction. The product was extract with DCM and wash with brine and dry by $Na_2SO_4$. The crude residue was purified by silica gel chromatography (0-10% MeOH in DCM) to afford 14 mg (79%) of TH3-189 as a yellow-white oil. $^1H$ NMR (500 MHz, $CDCl_3$) δ 7.46 (d, *J* = 1.9 Hz, 1H), 7.41 (dd, *J* = 8.0, 1.9 Hz, 1H), 7.06 (d, *J* = 8.0 Hz, 1H), 6.40 (dd, *J* = 16.8, 1.9 Hz, 1H), 6.07 (dd, *J* = 16.7, 10.3 Hz, 1H), 5.57 (dd, *J* = 10.4, 1.9 Hz, 1H), 4.92 (dd, *J* = 13.2, 1.9 Hz, 1H), 3.79 (d, *J* = 13.6 Hz, 1H), 3.59 (d, *J* = 13.5 Hz, 1H), 3.14 (dt, *J* = 8.8, 4.3 Hz, 1H), 3.10 (s, 1H), 2.68

(ddd, *J* = 32.9, 15.7, 9.9 Hz, 2H), 2.54 (q, *J* = 8.8 Hz, 1H), 2.03 (tdd, *J* = 11.5, 8.1, 5.1 Hz, 1H), 1.90 – 1.75 (m, 2H), 1.50 (dtd, *J* = 19.7, 9.9, 6.3 Hz, 1H). $^{13}$C NMR (126 MHz, $CDCl_3$) δ 164.68, 142.56, 134.59, 132.35, 130.24, 128.83, 128.49, 128.10, 122.18, 82.61, 78.54, 66.53, 56.64, 54.58, 49.75, 28.50, 21.20. HRMS (ESI) m/z calcd for $C_{17}H_{19}N2O^+$ $[M+H]^+$ : 267.1489; found: 267.1484

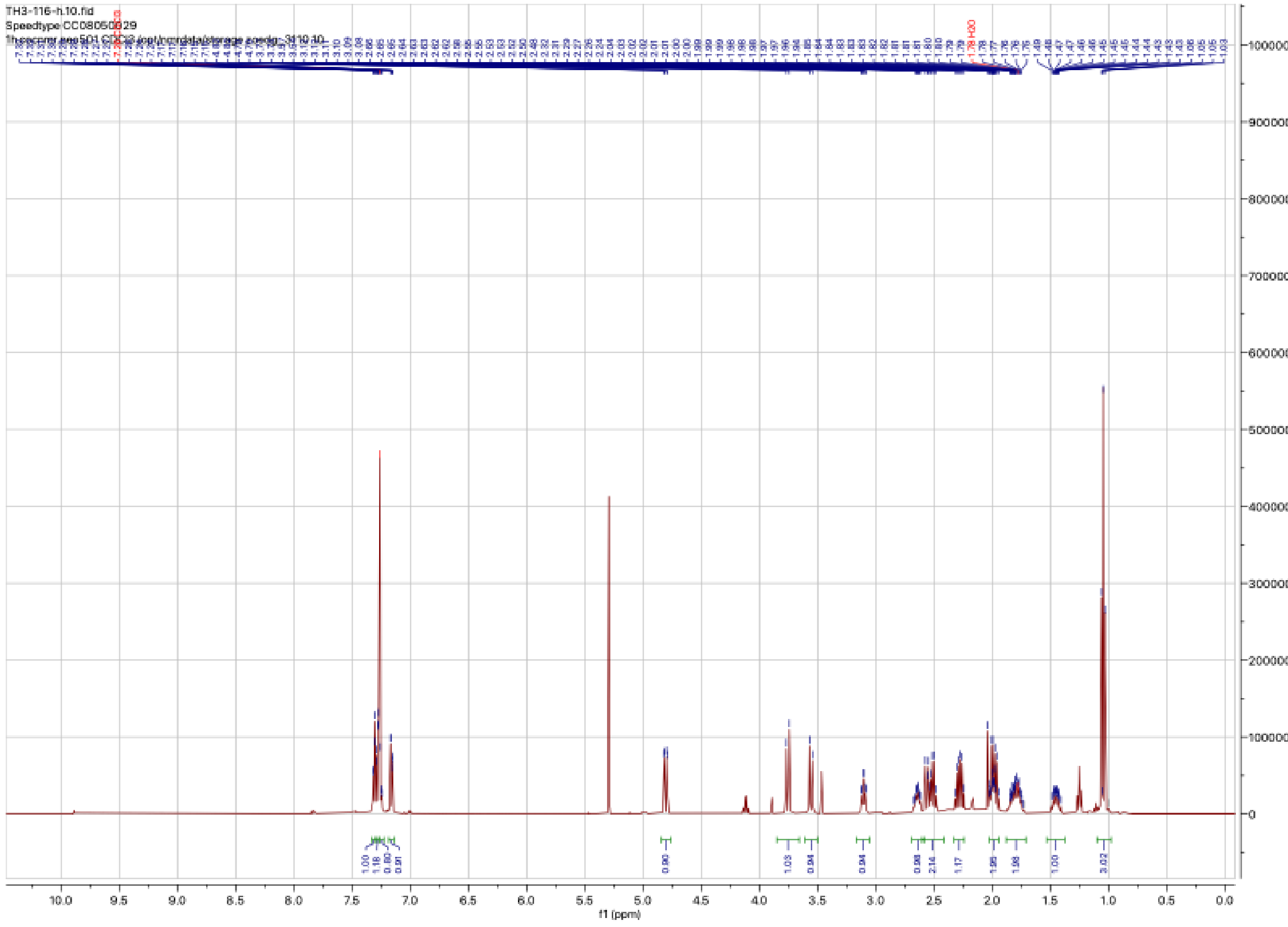


$^{1}$H NMR spectra of TH3-116

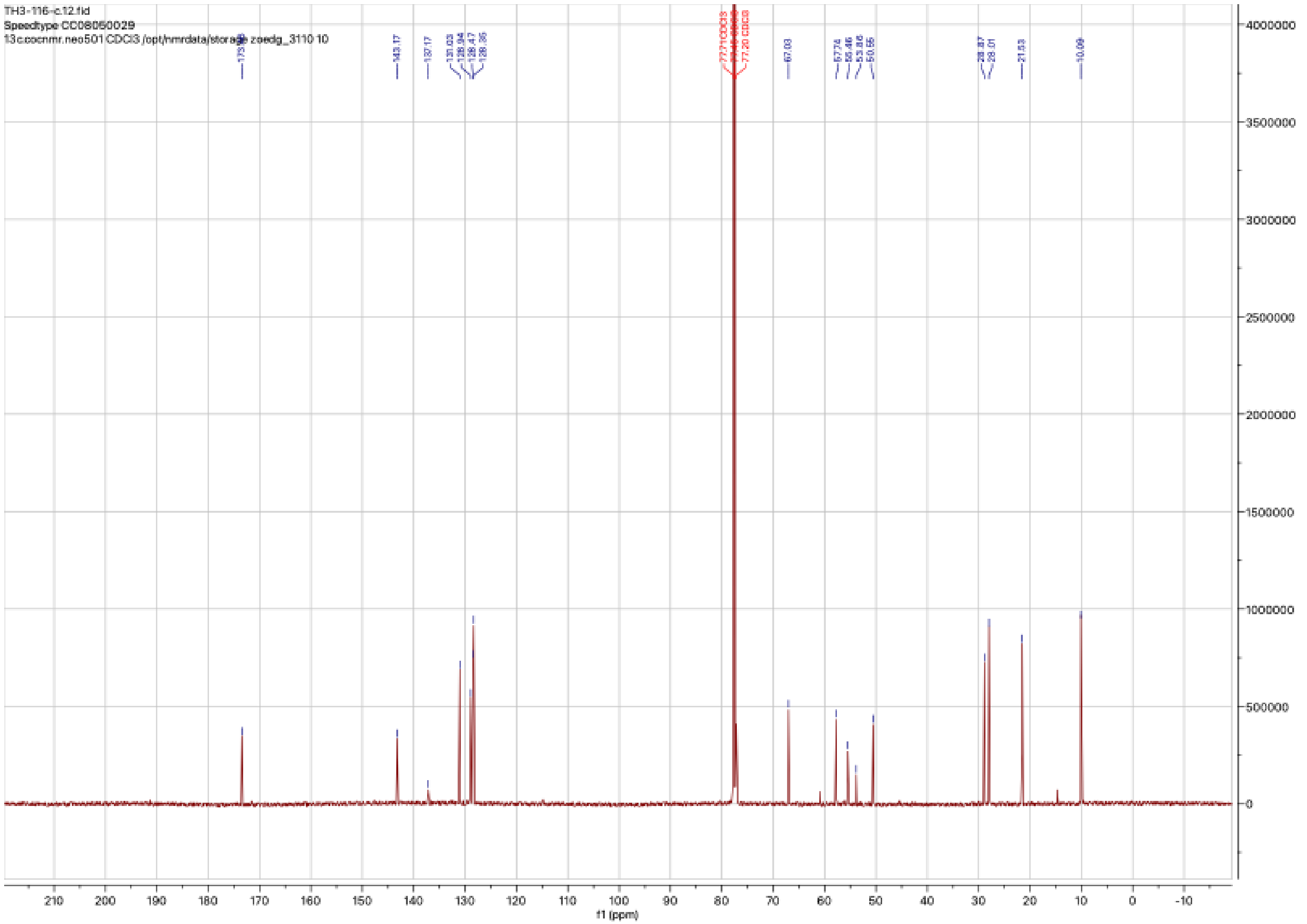


$^{13}C$ NMR spectra of TH3-116

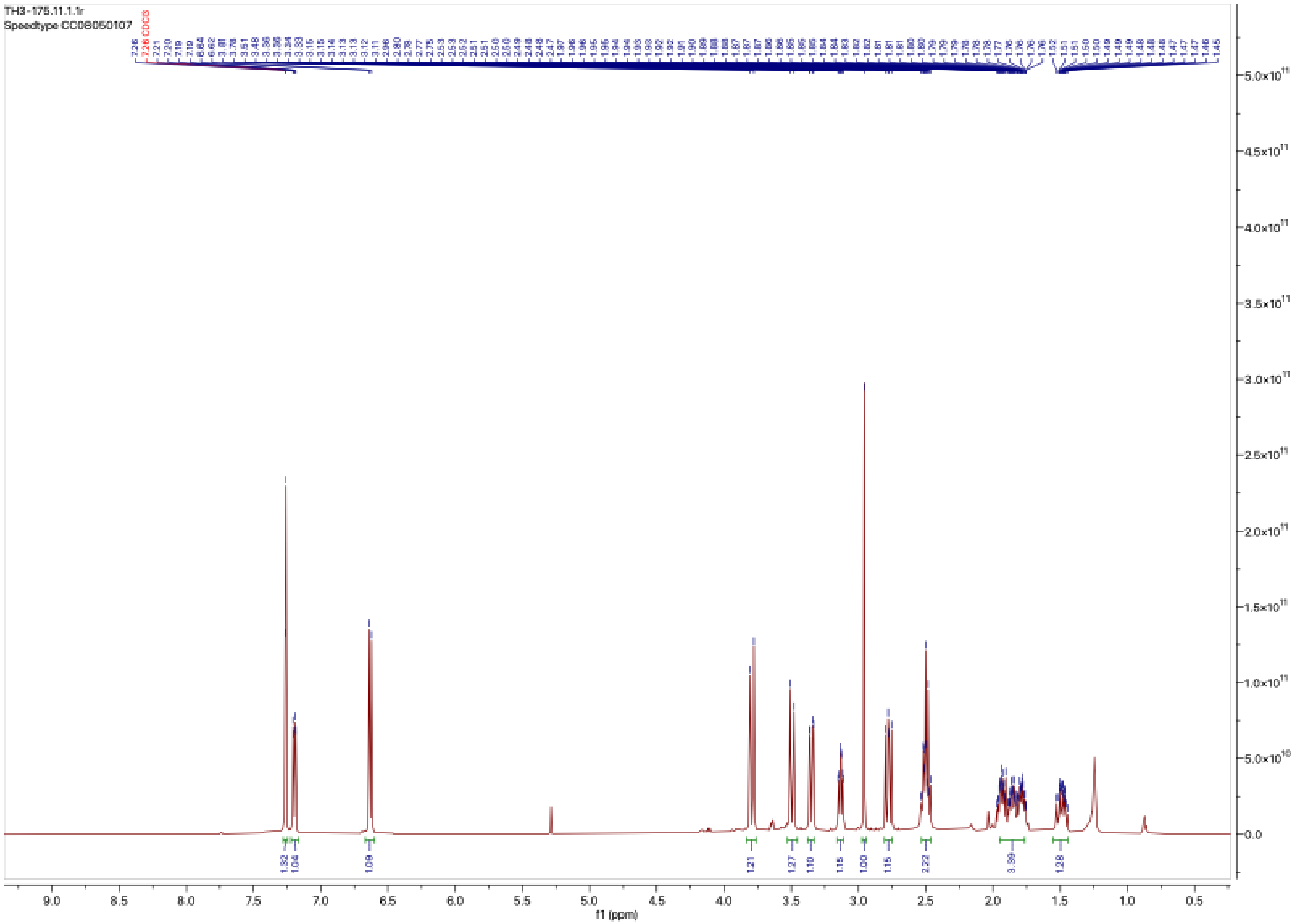

$^1H$ NMR spectra of TH3-175

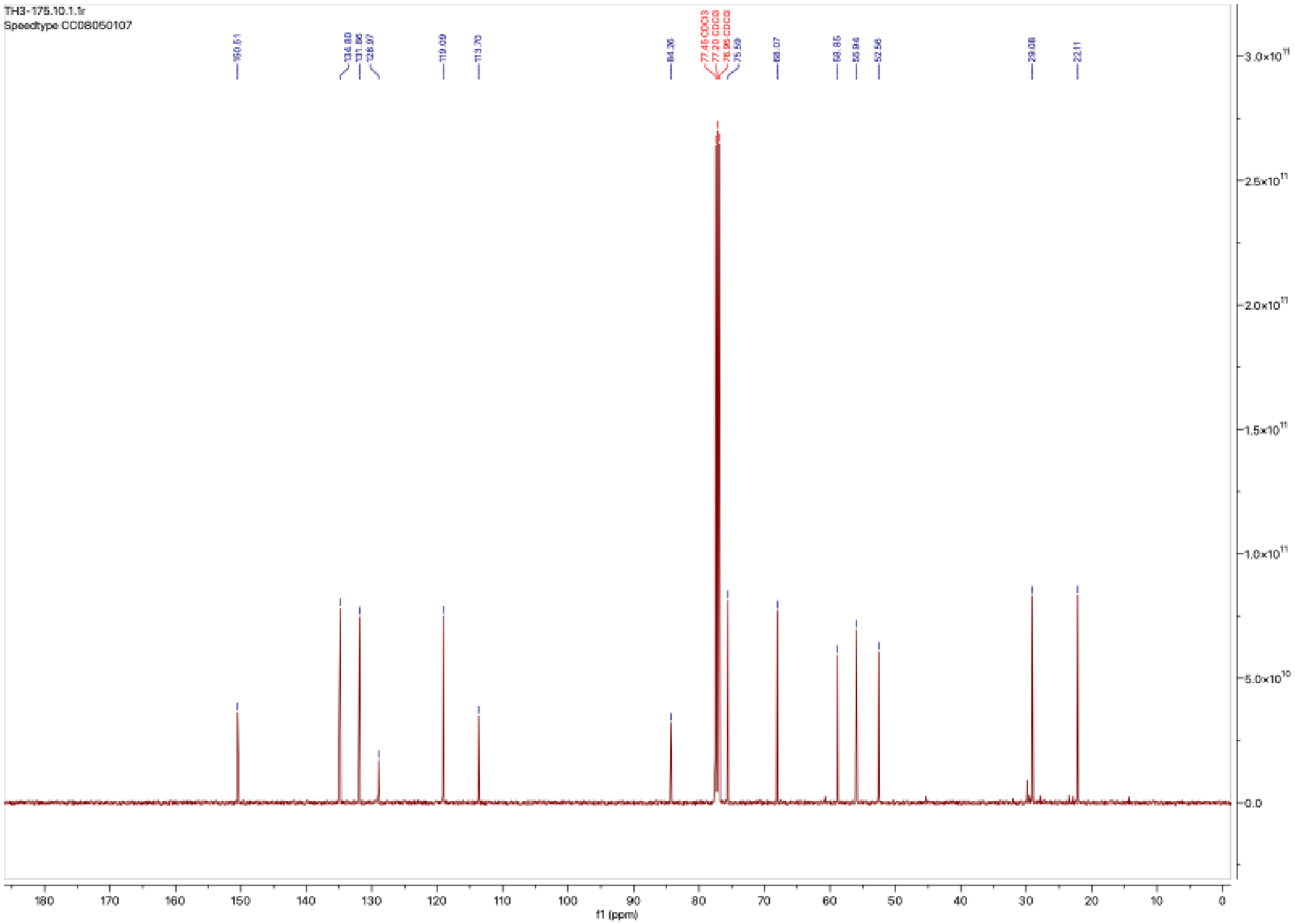


$^{13}C$ NMR spectra of TH3-175

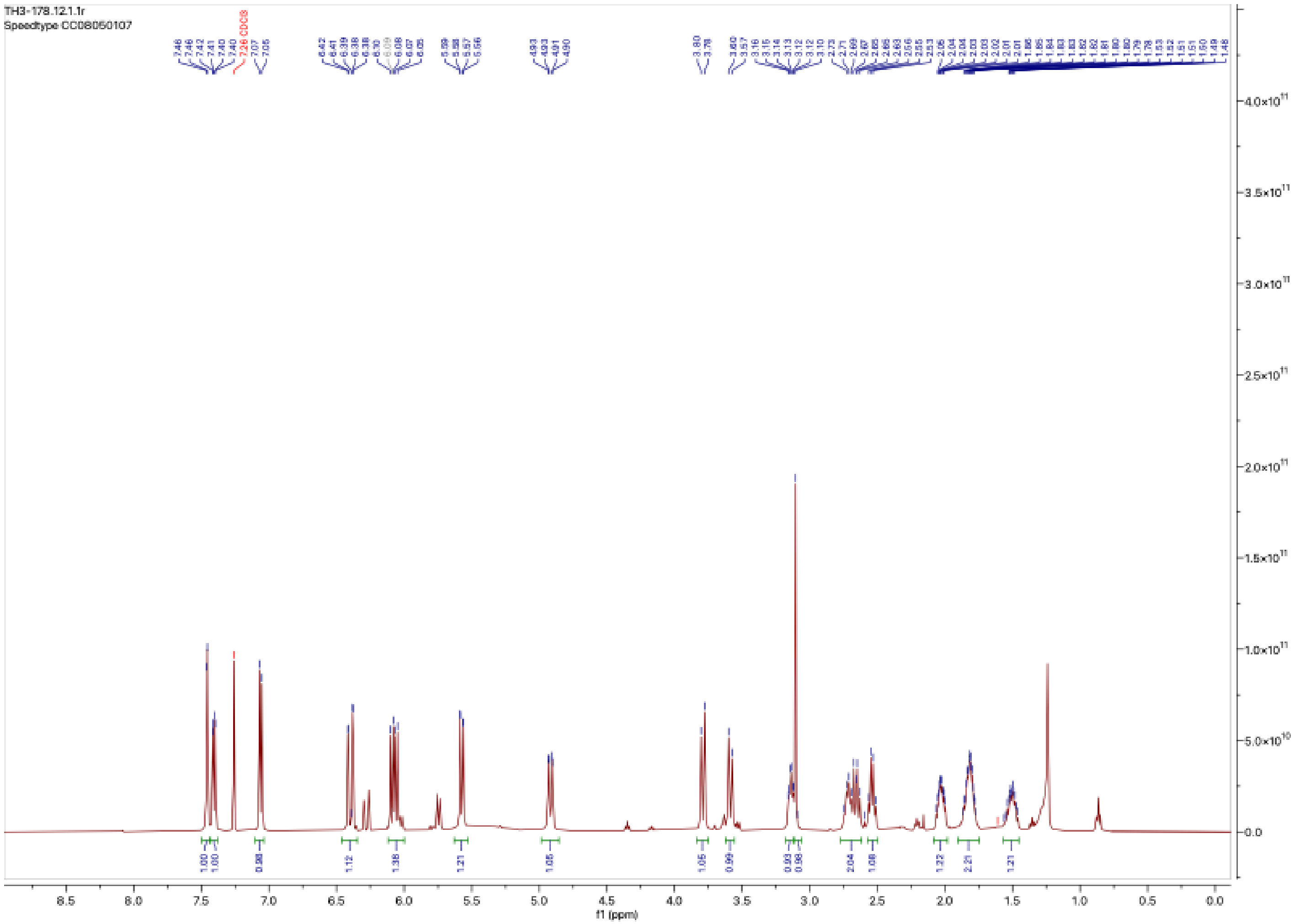


[1]H NMR spectra of TH3-189

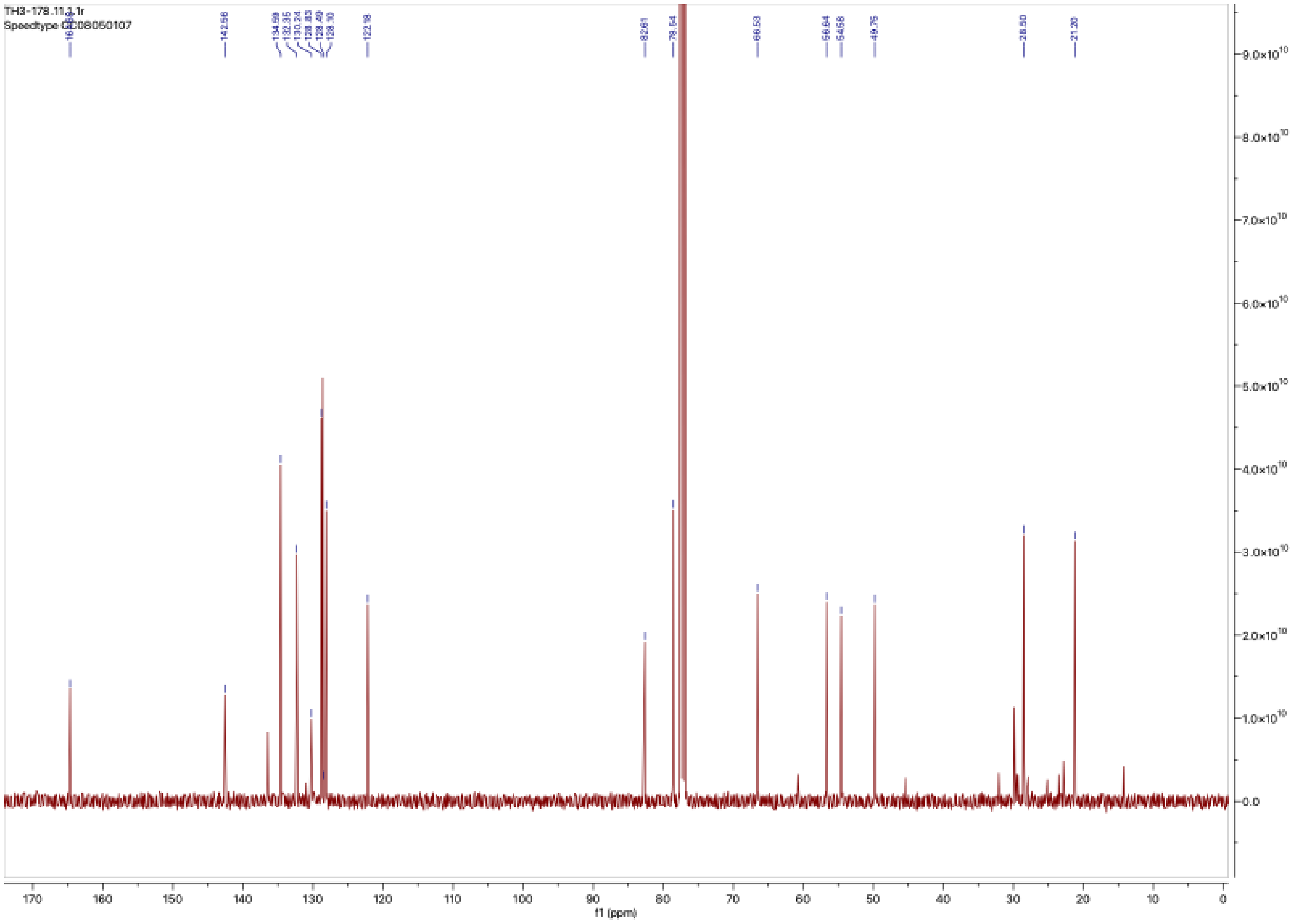


$^{13}C$ NMR spectra of TH3-189
